# Inhibition of microbial deconjugation of micellar bile acids protects against intestinal permeability and liver injury

**DOI:** 10.1101/2021.03.24.436896

**Authors:** Darrick K. Li, Snehal N. Chaudhari, Mozhdeh Sojoodi, Yoojin Lee, Arijit A. Adhikari, Lawrence Zukerberg, Stuti Shroff, Stephen Cole Barrett, Kenneth Tanabe, Raymond T. Chung, A. Sloan Devlin

**Affiliations:** Liver Center, Massachusetts General Hospital, Harvard Medical School, Boston, MA, USA; Department of Biological Chemistry and Molecular Pharmacology, Blavatnik Institute, Harvard Medical School, Boston, MA, USA; Department of Surgery, Massachusetts General Hospital, Harvard Medical School, Boston, MA, USA; Department of Pathology, Massachusetts General Hospital, Harvard Medical School, Boston, MA, USA

## Abstract

Altered host-microbe interactions and increased intestinal permeability have been implicated in the pathogenesis of a range of diseases. However, the mechanisms by which gut microbes affect epithelial barrier integrity remain unclear. Few host-produced metabolites that protect against epithelial damage have been identified, and whether microbial metabolism of host factors alters intestinal barrier function is unknown. Here, we investigate the effects of bacterial metabolism of host-produced bile acid (BA) metabolites on epithelial barrier integrity. We observe that rats fed a choline-deficient, high-fat diet (CDAHFD) exhibit reduced abundance of host-produced conjugated BAs in the intestine at early timepoints coinciding with increased permeability. We show that in vitro, conjugated BAs protect gut epithelial monolayers from damage caused by bacterially produced unconjugated BAs through micelle formation. We then demonstrate that inhibition of BA deconjugation using a small molecule inhibitor of gut bacterial bile salt hydrolase (BSH) enzymes prevents development of pathologic intestinal permeability and hepatic inflammation in CDAHFD-fed rats. Finally, we show that the predominant conjugated BAs in humans protect against epithelial barrier disruption in vitro. Our study identifies a protective role for conjugated BAs in intestinal epithelial barrier function and suggests that rational manipulation of microbial BA metabolism could be leveraged to regulate gut barrier integrity.

## Introduction

In health, the intestinal epithelium forms a dynamic and tightly sealed barrier that is selectively permeable (Buckley & Turner, 2018). However, under pathologic conditions, tight junctions can become disrupted with excessive leakage of dietary and bacterial antigens, including lipopolysaccharide (LPS), into the portal and systemic circulation, directly inducing inflammation in extraintestinal organs (Massier, Bluher, Kovacs, & Chakaroun, 2021). Emerging data have implicated increased intestinal permeability in the pathogenesis of a range of human diseases, including inflammatory bowel disease (IBD), liver disease, type 1 and type 2 diabetes, cardiovascular disease, and depression (Bosi et al., 2006; Chopyk & Grakoui, 2020; Damms-Machado et al., 2017; Ohlsson et al., 2019; Pasini et al., 2016). The gut microbial communities of these patients are altered compared to healthy subjects, and gut microbial imbalance has been proposed to contribute to the development of intestinal permeability (Albillos, de Gottardi, & Rescigno, 2020; Chakaroun, Massier, & Kovacs, 2020). However, specific host and microbial factors that result in the development of pathologic intestinal permeability remain unclear.

Bile acids (BAs) have been implicated as potential causal agents in the development of pathogenic intestinal permeability (Murakami, Tanabe, & Suzuki, 2016; Papillon, Frey, Ford, & Gayer, 2013). BAs are steroidal natural products that are present in high concentrations in the gut (∼200 μM to 1 mM in the large intestine) (Hamilton et al., 2007). In the liver, host enzymes convert cholesterol into conjugated primary bile acids, which contain a steroidal core appended to taurine or glycine through an amide bond linkage (**Figure 1A**). These compounds act as digestive surfactants; in humans, these metabolites are stored in the gallbladder and secreted into the small intestine post-prandially in order to aid in the absorption of lipids and vitamins (de Aguiar Vallim, Tarling, & Edwards, 2013). In the lower GI tract, bacteria convert primary conjugated bile acids into unconjugated primary BAs through the action of bile salt hydrolase (BSH) enzymes (Ridlon, Harris, Bhowmik, Kang, & Hylemon, 2016). BSHs are expressed in a broad range of human gut bacteria (Song et al., 2019) and have no mammalian homolog (Foley, O’Flaherty, Barrangou, & Theriot, 2019). Bacterial enzymes then further chemically modify primary BAs, producing unconjugated secondary BAs. Following enterohepatic recirculation, these molecules can be converted to conjugated primary and secondary BAs in the liver and then re-secreted into the gut (Ridlon et al., 2016).

**Figure 1.**
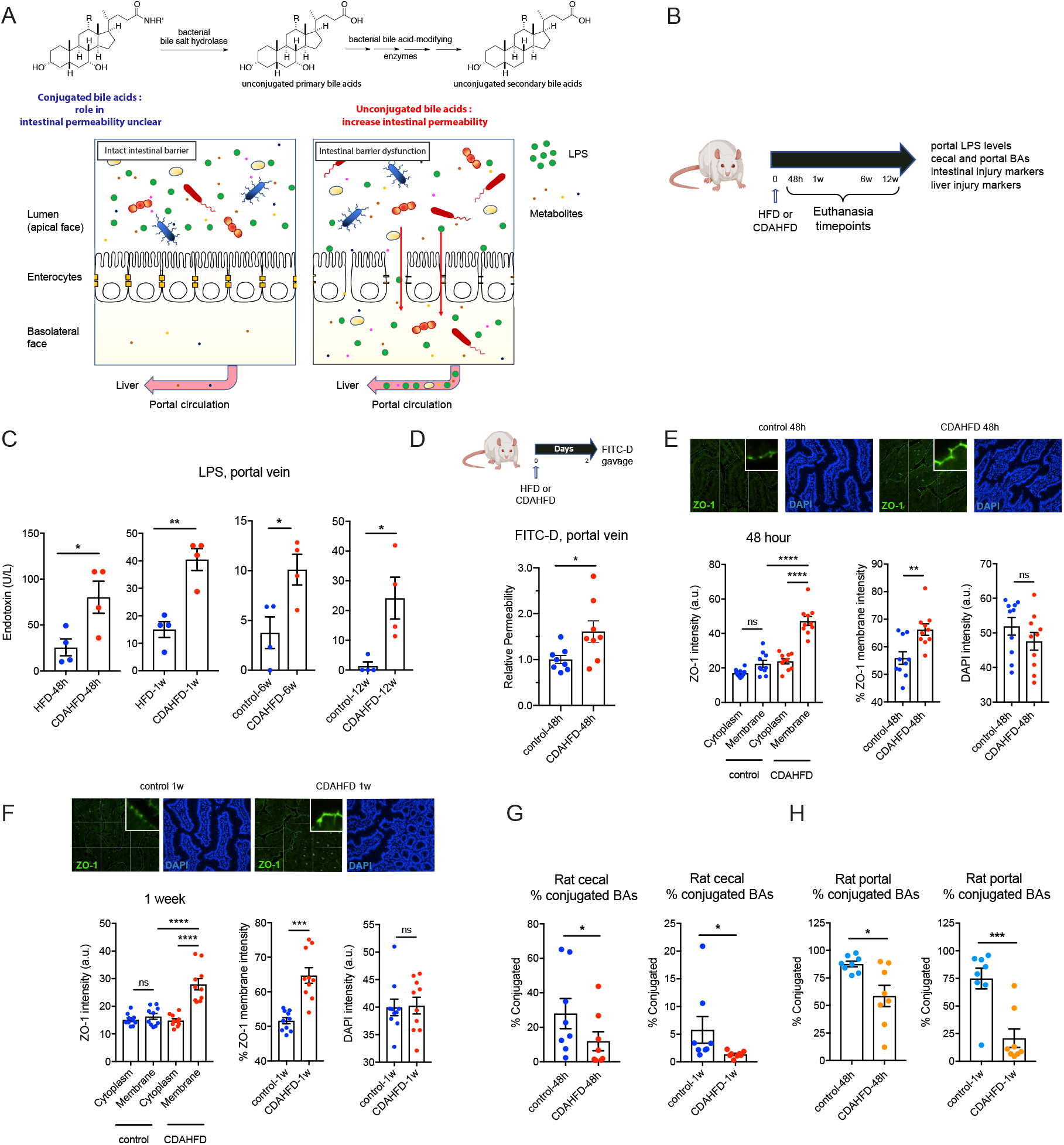
CDAHFD-fed rats develop increased intestinal permeability at early timepoints coinciding with decreased conjugated BA abundance. (A) Human gut bacterial enzymes transform conjugated primary BAs into unconjugated secondary BAs. While hydrophilic unconjugated BAs damage epithelial integrity, the roles of unconjugated BAs and bacterial BSHs in intestinal membrane integrity are unclear. (B) Schematic of rat intestinal permeability timecourse experiment. A choline-deficient, L-amino acid defined, high-fat diet was fed to rats to induce intestinal permeability and liver damage. High fat diet-fed rats served as controls. Rats were euthanized at 48 hours, 1 week, 6 week, or 12 week timepoints. Tissues and blood were collected for metabolite quantification and evaluation of intestinal and liver injury markers. (C) Portal venous levels of lipopolysaccharide (LPS) were significantly increased in CDAHFD-fed rats compared to control rats after 48h, 1w, 6w, and 12w on diet (n=4 per group, two-tailed Welch’s t test). (D) CDAHFD-fed rats developed increased intestinal permeability after 48h of diet. Measurement of FITC-Dextran levels in systemic circulation after gavage in control and CDAHFD-fed rats (n=8 per group, two-tailed Welch’s t test). (E-F) CDAHFD induced increased expression and apical membrane localization of ZO-1 at early timepoints. ZO-1 immunofluorescence staining of ileum from control and CDAHFD-fed rats at 48h (E) and 1w (F) timepoints (n=10 intestinal cells quantified per group, see Methods for statistical analyses). (G, H) Conjugated BA abundance was reduced at early timepoints (48h, 1w) in the cecum (G) and portal vein (H) of CDAHFD-fed rats as determined by UPLC-MS BA analysis from control and CDAHFD-fed animals (n=8 per group, two-tailed Welch’s t test, see Supplementary Information for concentrations of total and individual BAs in cecal contents and portal vein at 48h, 1w). See Methods for statistical analyses. \**p*<0.05, \*\**p*<0.005, \*\*\**p*<0.001, \*\*\*\**p*<0.0001. Bars represent mean ± SEM.

Previous work has demonstrated that exposure of epithelial monolayers to certain hydrophobic BAs, including unconjugated BAs, leads to increased intestinal permeability in vitro and may contribute to the development of intestinal inflammation and disruption of intestinal homeostasis in vivo (Raimondi et al., 2008; Sarathy et al., 2017). In animal models, long-term administration of a high-fat diet has been shown to lead to increased intestinal permeability associated with an enrichment of hydrophobic BAs in the intestinal BA pool (Gupta et al., 2020; Stenman, Holma, Eggert, & Korpela, 2013; Stenman, Holma, & Korpela, 2012). While unconjugated BAs have been shown to increase epithelial permeability, the role of conjugated BAs in epithelial permeability is unclear, and it is not known whether bacterial BA deconjugation affects intestinal homeostasis in vivo.

Considering the close physiological relationship between the gut and the liver, optimal intestinal barrier function is crucial for liver homeostasis (Nicoletti et al., 2019). Disruption of intestinal barrier integrity may accelerate the pathogenesis of liver disease (Chopyk & Grakoui, 2020). Here, we observe that increased intestinal permeability is an early feature of liver disease in rats fed a choline-deficient, L-amino acid defined, high-fat diet, an animal model of liver cirrhosis accompanied by pathogenic intestinal barrier damage (Longo et al., 2020). We also observed a significant reduction in the abundance of intestinal and portal venous conjugated BAs in CDAHFD-treated rats at early timepoints. Hypothesizing that conjugated bile acids could protect the intestinal barrier against chemical insults, we demonstrate that tauro-conjugated BAs sequester unconjugated BAs into micelles. We then show that this sequestration of unconjugated bile acids protects epithelial cells from damage and permeability in vitro. Furthermore, using a small molecule inhibitor of bacterial BSHs that we recently developed (Adhikari et al., 2021; Adhikari et al., 2020), we show that inhibition of BSH activity increases conjugated BA abundance in vivo and prevents the development of increased intestinal permeability, hepatic steatosis, and hepatic inflammation in diseased rats. Finally, we demonstrate that glyco-conjugated BAs, the predominant BAs in humans, also protect against epithelial damage caused by unconjugated BAs in vitro. Together, these data indicate that gut microbial enzymes interact with host metabolites to affect intestinal barrier function.

## Results

### CDAHFD-fed rats develop increased intestinal permeability prior to developing hepatic inflammation

To investigate the molecular mechanisms leading to the development of intestinal permeability in the context of an animal model relevant to human disease, we utilized a choline-deficient, L-amino acid defined, high-fat diet (CDAHFD) in rats. CDAHFD is an established rodent model of diet-induced cirrhosis that is characterized by disrupted epithelial barrier integrity (Zhong, Zhou, Xu, & Gao, 2020). Previous studies have also found choline deficiency to be associated with development of non-alcoholic fatty liver disease (NAFLD) and NASH in humans (Corbin & Zeisel, 2012; Guerrerio et al., 2012). CDAHFD-fed rats developed cirrhosis in 12 weeks (Matsumoto et al., 2013) while controls fed a high-fat diet with equivalent fat by weight developed microvesicular steatosis without inflammation or fibrosis (**Figure S1A**). CDAHFD-fed rats gained less weight than controls and exhibited significantly increased markers of hepatocellular injury, hepatic hydroxyproline levels, and mRNA expression of fibrosis-related and inflammatory genes at 12 weeks post-diet intervention (**Figure S1B-C**, refer **Table S1** for all qPCR primer sequences).

To assess intestinal permeability, we measured portal lipopolysaccharide (LPS) in CDAHFD-fed rats at 48 hours, 1 week, 6 weeks, and 12 weeks (**Figure 1A,B**). Remarkably, portal LPS levels were higher in CDAHFD-fed rats compared to controls as early as 48 hours post-diet intervention (**Figure 1C**), when markers of liver injury were not yet significantly elevated (**Figure S1C-E**). Further, intestinal epithelium from CDAHFD-fed rats exhibited significantly increased inflammation and epithelial hyperplasia at early time points (**Figure S1F**). These data suggest that intestinal injury and permeability precede hepatic phenotypes in the CDAHFD model of liver disease. To confirm these results, using a FITC-Dextran assay, we found that CDAHFD-fed rats exhibited a significant increase in intestinal permeability compared to controls at 48 hours after diet initiation (**Figure 1D**). We observed significantly increased membrane localization of ZO-1 in intestinal epithelial cells of CDAHFD-fed rats at 48 hours that persisted after 1 week of diet (**Figure 1E, F**). Increased expression and plasma membrane localization of ZO-1 are indicative of increased intestinal permeability (Guo, Wang, Liu, & Ye, 2018; Karczewski et al., 2010). Moreover, treatment of gut epithelial cells with permeability agents results in dynamic changes in ZO-1 subcellular localization, including recruitment to the plasma membrane (Guan et al., 2011; Odenwald et al., 2017). As such, ZO-1 redistribution in the intestinal epithelium of CDAHFD-fed rats is consistent with increased intestinal permeability. Importantly, at 48 hours, CDAHFD-fed rats did not exhibit any evidence of hepatic inflammation compared to controls while inflammation was apparent at 1 week post-dietary intervention (**Figure S1C-E**). Together, these results demonstrate that intestinal permeability is an early feature of this animal model and precedes the development of hepatic inflammation.

### Conjugated BAs are reduced in the cecum and portal veins of CDAHFD-fed rats at early timepoints

To assess whether changes in BAs are associated with increased intestinal permeability, we performed intestinal and portal venous BA profiling in CDAHFD-fed and control rats using ultra-high performance liquid chromatography-mass spectrometry (UPLC-MS). We observed a significant decrease in the abundance of cecal and portal venous conjugated BAs in CDAHFD-fed rats at 48 hours and 1 week after dietary intervention (**Figure 1G-H**). In particular, levels of tauro-cholic acid (TCA) and tauro-alpha- and tauro-beta-muricholic acid (Tα/βMCA) were decreased in both cecal contents and portal veins at both timepoints (**Figure S2-5)**. Total cecal and portal venous BA concentrations were similar between the two groups at these early timepoints (**Figure S2-5**). In addition, levels of the conjugated bile acids tauro-chenodeoxycholic acid (TCDCA), tauro-deoxycholic acid (TDCA), and tauro-ursodeoxycholic acid (TUDCA) were decreased and levels of the unconjugated BAs cholic acid (CA), chenodeoxycholic acid (CDCA), α-muricholic acid (αMCA), and β-muricholic acid (βMCA) were increased in portal veins at the 1 week timepoint (**Figure S5**). These findings indicate that changes in cecal and portal venous BA composition are associated with the onset of increased intestinal permeability in CDAHFD-fed rats and suggest that these metabolites could play a causal role in development of gut barrier damage.

Intestinal permeability and inflammation are regulated in part by the BA-sensing farnesoid X receptor (FXR), which has been linked to NAFLD/NASH pathogenesis (Arab, Karpen, Dawson, Arrese, & Trauner, 2017; Li et al., 2013). In addition, FXR signaling modulates a tight negative feedback loop that controls BA synthesis in the liver (Fiorucci & Distrutti, 2015). At 48 hours after dietary intervention, ileal *Fgf15* expression was similar in both groups when multiple lines of evidence for increased intestinal permeability were already observed in CDAHFD-fed animals (**Figure S6A)**. Expression of BA synthesis and conjugation enzymes were not significantly affected in CDAHFD-fed rats at 48 hours of diet **(Figure S6B)**. Expression levels of BA transporter proteins were also unaffected in CDAHFD-fed rats compared to controls **(Figure S6C)**. We found that cecal BSH activity in CDAHFD rats was significantly increased at 48 hours compared to control rats (**Figure S6D**) (Adhikari et al., 2020). These data suggest that the decrease in conjugated BAs in CDAHFD-fed rats was driven at least in part by an increase in BA deconjugation by gut microbes. Together, our findings suggest that FXR-independent mechanisms are responsible for the earliest events that establish intestinal barrier dysfunction in this animal model of pathogenic intestinal permeability.

### In vivo BA compositions reflect increased intestinal permeability as disease progresses

UPLC-MS analysis revealed significant decreases in cecal unconjugated BAs in CDAHFD-fed rats at 6 weeks (e.g., αMCA, LCA, DCA, UDCA) and in unconjugated, conjugated, and total BAs at 12 weeks (e.g., CA, αMCA, βMCA, LCA, DCA, UDCA, TDCA, tauro-omega-muricholic acid (TωMCA), TUDCA) compared to control animals (**Figure S7**,**8)**. Interestingly, at the same timepoints, portal venous BA profiling revealed the inverse finding, with significantly higher concentrations of total and unconjugated BAs (e.g., CA, CDCA, αMCA, βMCA) in CDAHFD-fed rats compared to controls (**Figure S9**,**10)**. No significant differences were observed in the expression of genes involved in BA synthesis (*Cyp7a1, Cyp8b1, Cyp27a1*) or transport (*Asbt, Osta/Ostb*) (**Figure S11A**,**B**), indicating that the observed changes are unlikely to be the result of decreased BA synthesis or increased BA transport. These findings are consistent with the observation that CDAHFD-treated rats exhibit epithelial barrier damage and suggest that as disease progresses, increased intestinal permeability results in leakage of intestinal BAs into portal circulation.

### Conjugated BAs protect intestinal epithelial monolayers from unconjugated BA-mediated damage in vitro

Based on our in vivo observations, we hypothesized that groups of conjugated or unconjugated BAs could be causally contributing to epithelial layer damage. To test this hypothesis, we utilized Caco-2 cells in transwell inserts as an in vitro model system. Once differentiated, these cells form a polarized monolayer that mimics the intestinal epithelium, and this system has been previously used to study the effect of metabolites on gut layer integrity (Scott, Fu, & Chang, 2020). Gut permeability was assayed by measuring FITC-Dextran (4 kDa) permeability through the monolayer and quantifying fluorescence in the basolateral chamber (Chaudhari, Harris, et al., 2021). We found that cecal extracts isolated from CDAHFD-fed rats at 48 hours and 1 week induced increased permeability compared to cecal extracts from control rats (**Figure 2A**). These findings demonstrate that intestinal contents from CDAHFD-fed rats induce epithelial barrier permeability at early timepoints.

**Figure 2.**
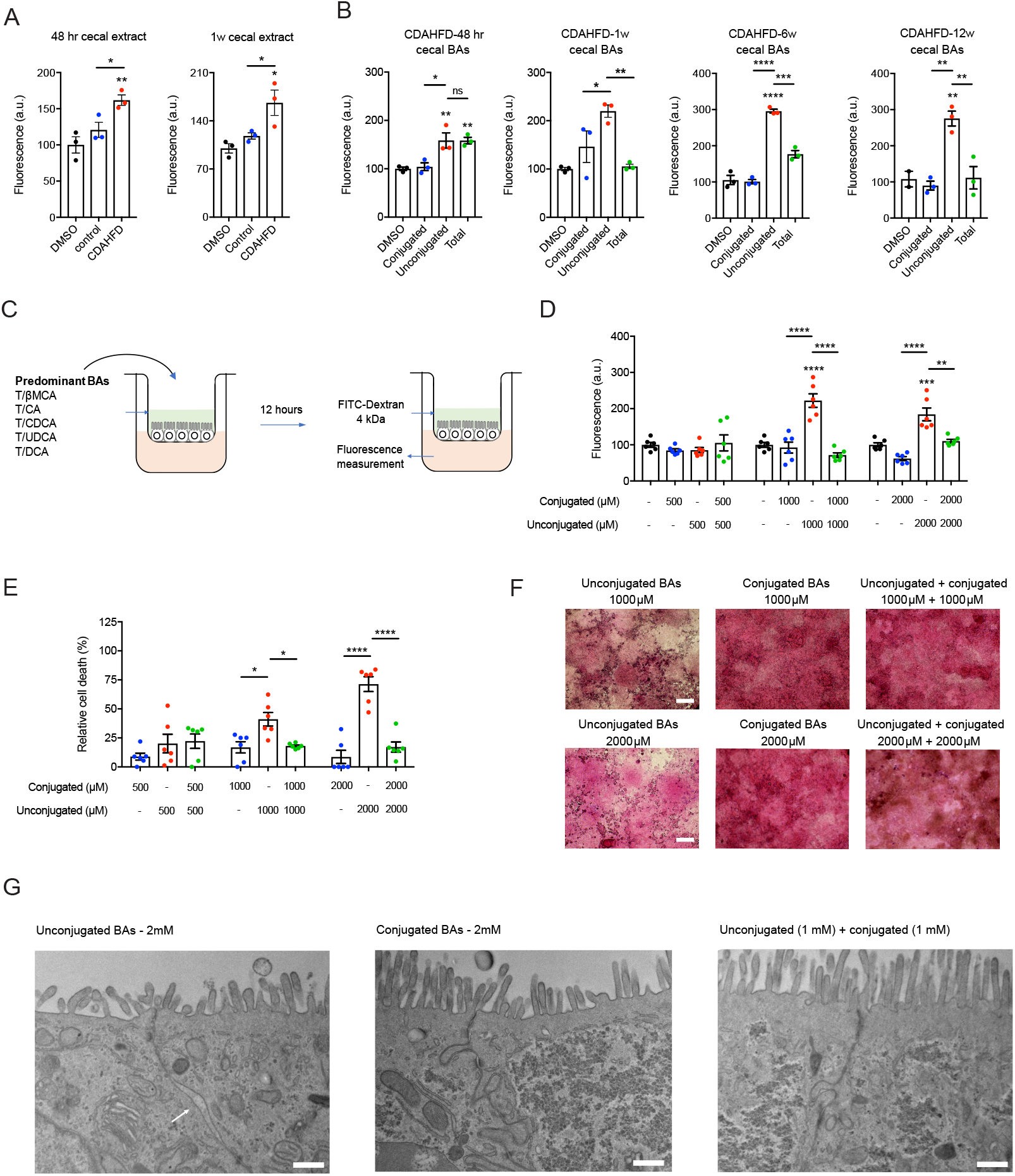
Conjugated BAs protect epithelial monolayers from unconjugated BA-induced permeability. (A) Cecal extracts from CDAHFD-fed rats at early timepoints caused increased epithelial permeability. Caco-2 monolayer permeability after exposure to purified cecal extracts from control and CDAHFD-fed rats at indicated timepoints as measured by FITC-Dextran passage into the basolateral chamber over 12 hours (n=3 per group, see Methods for statistical analyses). (B) Conjugated BAs alone did not increase Caco-2 monolayer permeability at concentrations found in the cecum of CDAHFD rats, and when combined with unconjugated BAs, conjugated BAs protected against unconjugated BA-induced permeability (n=3 per group, see Methods for statistical analyses). (C) Schematic of in vitro permeability experiment. Conjugated BAs (equimolar concentrations of TβMCA, TCA, TCDCA, TUDCA, TDCA), unconjugated BAs (equimolar concentrations of βMCA, CA, CDCA, UDCA, DCA), and combined BA pools (equimolar concentrations of both pools) were added to the apical chamber of a transwell system containing a Caco-2 monolayer and incubated for 12 hours before addition of FITC-Dextran 4kDa and fluorescence measurement from basolateral chamber. (D) Conjugated BAs protected against unconjugated BA-induced Caco-2 monolayer permeability at physiologic concentrations (n=6 per group, see Methods for statistical analyses). (E) Conjugated BAs protected epithelial monolayers from unconjugated BA-induced cell death. Cell viability of Caco-2 cells measured by MTT assay (n=6 per group, see Methods for statistical analyses). (F) Conjugated BAs protected the physical integrity of epithelial monolayers from unconjugated BA-induced damage. Representative light microscopy images of H&E-stained Caco-2 monolayers after exposure to BA pools. Scale bar=20 μm. (G) Conjugated BAs prevented the development of unconjugated BA-induced tight junction dilatation. Representative TEM images of Caco-2 cells from transwells after exposure to BA pools at indicated concentrations. The white arrow points to tight junction dilatation. Scale bar=500 nm. see Supplementary Information for additional representative TEM images. Unless otherwise specified, all experiments were performed in triplicate. See Methods for statistical analyses. Data not marked with asterisk(s) are not significant. \**p*<0.05, \*\**p*<0.005, \*\*\**p*<0.001, \*\*\*\**p*<0.0001. Bars represent mean ± SEM.

To test whether cecal BAs specifically induce epithelial permeability, we generated reconstituted pools of BAs that mimic the average physiological concentrations observed in rat ceca and tested their ability to induce permeability in Caco-2 monolayers in vitro. The 48 hour CDAHFD total BA pool induced significantly higher permeability than the DMSO control (**Figure 2B**). Interestingly, while the unconjugated BA pool alone induced a significant increase in permeability, the conjugated BA pool did not damage the monolayer barrier integrity. Remarkably, at later time points (1, 6, 12 weeks), the addition of conjugated BAs to unconjugated BAs mitigated the severity of epithelial permeability compared to unconjugated BAs alone (**Figure 2B**).

We next hypothesized that cecal conjugated BAs protect against the disruption of the epithelial integrity. To test this hypothesis, we generated equimolar pools of the predominant cecal BAs that we detected in vivo. We treated differentiated Caco-2 monolayers with increasing concentrations of either (1) unconjugated BAs (βMCA, CA, DCA, UDCA, CDCA); (2) conjugated BAs (TβMCA, TCA, TDCA, TUDCA, TCDCA) or (3) combined BA pools, followed by the permeability assessment (**Figure 2C**).

At physiological concentrations of BAs in CDAHFD-fed rat cecal contents (∼1000-4000 μM), epithelial monolayer integrity was compromised after addition of unconjugated BAs but damage was prevented by addition of an equimolar concentration of conjugated BAs (**Figure 2D, S2**,**3**,**7**,**8**). Conjugated BAs alone did not disrupt monolayer integrity at any concentrations tested. Similarly, while unconjugated BAs were toxic to cells, equimolar addition of conjugated BAs abrogated this effect (**Figure 2E**). Visualization of Caco-2 monolayers by hematoxylin and eosin staining further confirmed the toxic effects of unconjugated BAs were largely rescued by addition of conjugated BAs (**Figure 2F**). Finally, to assess the integrity of tight junctions in the epithelial monolayers, we performed transmission electron microscopy (TEM) on Caco-2 epithelial monolayers exposed to the above BA groups. We identified dilatations in the tight junctions of Caco-2 cells exposed to unconjugated BAs alone not present in cells exposed to conjugated BAs alone or to a combination of both (**Figure 2G, Figure S12**). These tight junction dilatations have been observed in ileal enterocytes of Crohn’s disease patients and correlate with increased permeability (Soderholm et al., 2002). Together, these results demonstrate that while unconjugated BAs damage epithelial layers, conjugated BAs protect against intestinal epithelial damage and permeability in vitro.

### Conjugated BAs sequester unconjugated BAs through the formation of micelles

We next sought to determine the mechanism by which conjugated BAs protect intestinal epithelial monolayers from unconjugated BA-mediated permeability in vitro. BAs are detergents and effectively solubilize fats and vitamins by forming micelles in the intestine (Hofmann, 1963). We hypothesized that when combined, conjugated BAs form micelles with unconjugated BAs, sequestering unconjugated BAs away from the epithelial cells. To test this hypothesis, we assessed the critical micelle concentration (CMC) of these BA populations using a fluorescent probe (Fluksman & Benny, 2019). Unconjugated BAs are expected to have a higher CMC compared to conjugated BAs (Pavlovic et al., 2018). If micelles formed after combining unconjugated and conjugated BA populations, we would expect the CMC of the combined populations to be lower than that seen with unconjugated BAs alone. Consistent with this hypothesis, the combined pool CMC was 4.2 mM while the unconjugated and conjugated BA pool CMCs were 6.7 mM and 4.0 mM, respectively (**Figure 3A**). The addition of 80 mM urea prevented micelle formation as previously described (**Figure 3A**) (Hofmann, 1963). We also performed direct visualization of micelles using negative staining-electron microscopy. At 5 mM concentration, no micelles were visualized in the unconjugated BA pool while micelles were seen in the conjugated BA pool, consistent with our CMC determinations and total cecal BA quantification in CDAHFD-fed rats at the 48 hour time point (**Figure 3B, S2**). Combining BA pools resulted in larger micelles, while urea prevented micelle formation entirely. Thus, in the presence of conjugated BAs, unconjugated BAs are sequestered in micelles.

**Figure 3.**
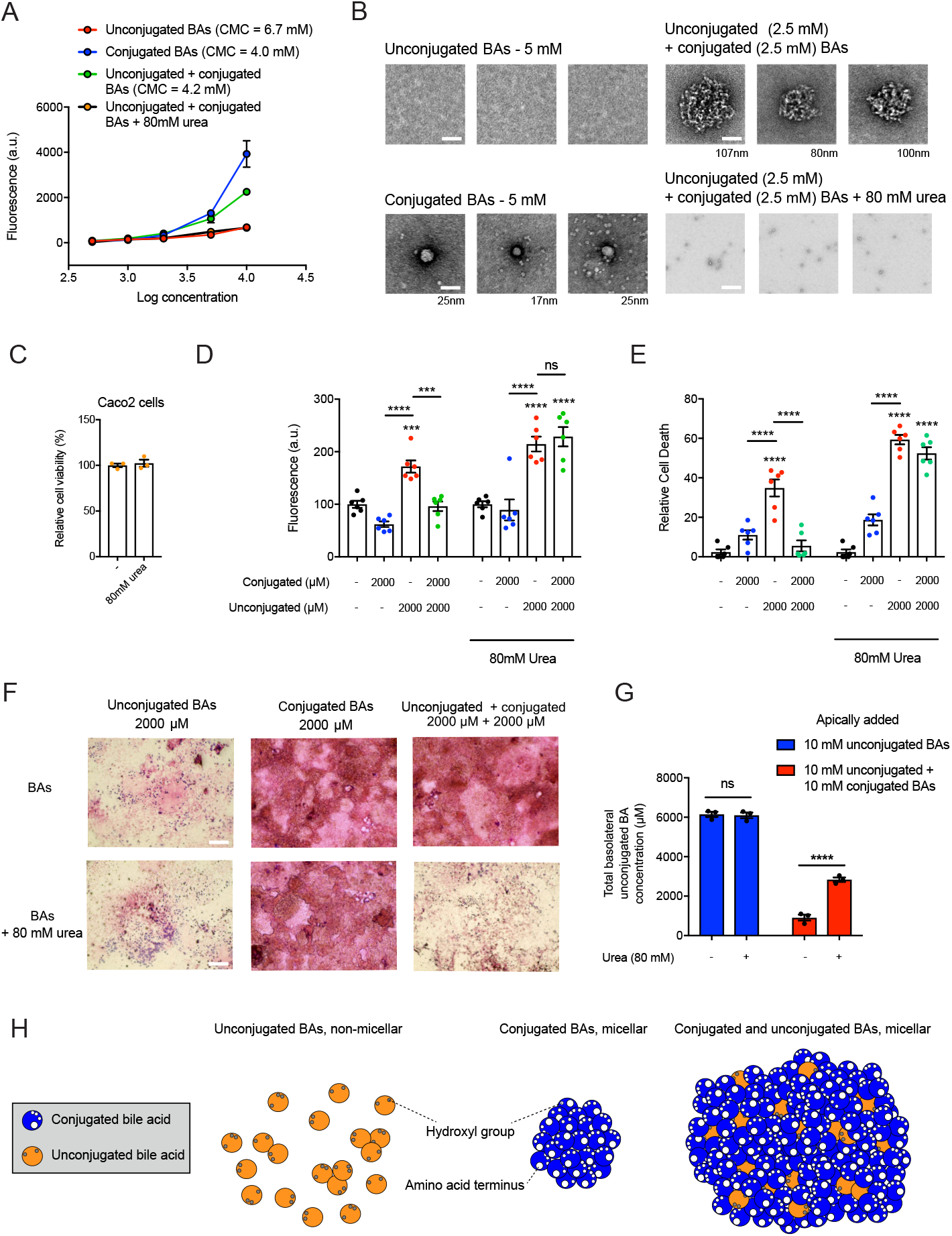
Conjugated and unconjugated BAs form micelles that sequester unconjugated BAs and prevent epithelial damage. (A) Conjugated BAs alone (equimolar concentrations of TβMCA, TCA, TCDCA, TUDCA, TDCA) and conjugated BAs combined with unconjugated BAs (equimolar concentrations of conjugated BAs with βMCA, CA, CDCA, UDCA, DCA) exhibited lower CMCs than unconjugated BAs alone. Micelle formation was disrupted after addition of urea (80 mM) (n=3 per group). (B) EM images of micelles formed from BA pools at indicated concentrations. While detectable micelles were visible in the conjugated BAs alone or unconjugated plus conjugated BA pools, no detectable micelles were visible in the unconjugated BAs or unconjugated plus conjugated BAs plus urea (80 mM) pools. Scale bar=50 nm, diameter of micelle indicated beneath each image (n=3 images per group). (C) MTT cell viability assay of Caco-2 cells in the presence or absence of 80 mM urea indicated that urea did not impact Caco-2 cell viability (n=3 per group, not significant by two-tailed Welch’s t test). (D) Micelle formation was necessary for the protective effect of conjugated BAs on epithelial permeability. Permeability measured by fluorescence (FITC-Dextran) in basolateral chamber in the presence or absence of 80 mM urea (n=6 per group, see Methods for statistical analyses). (E) Micelle formation was necessary for the protective effect of conjugated BAs on cell viability. Caco-2 cell viability measured by MTT assay in the presence or absence of urea (n=6 per group, see Methods for statistical analyses). (F) Micelle formation was necessary for the protective effect of conjugated BAs on epithelial layer integrity. Representative light microscopy images of H&E-stained Caco-2 monolayers in transwells in the presence or absence of urea. (G) Addition of urea to a mixed BA pool led to increased unconjugated BA passage across a Caco-2 monolayer. Quantification of basolateral concentrations of unconjugated BAs by UPLC-MS (n=3 per group, see Methods for statistical analyses). (H) Model for the sequestration of unconjugated BAs in micelles by conjugated BAs. BAs shown in cross-section and micellar shape depictions based on previous micelle models (Carey & Small, 1972; Faramarzi et al., 2017). While unconjugated BAs are largely non-micellar in solution, both conjugated BAs and mixtures of unconjugated and conjugated BAs form micelles. Unless otherwise specified, all experiments were performed in triplicate. See Methods for statistical analyses. Data not marked with asterisk(s) are not significant. \**p*<0.05, \*\**p*<0.005, \*\*\**p*<0.001, \*\*\*\**p*<0.0001. Bars represent mean ± SEM.

To investigate whether the protective effects of conjugated BAs are micelle-dependent, we tested whether BA pools induce intestinal permeability in the presence of 80 mM urea, a concentration that prevents micelle formation (**Figure 3A-B**) without inducing toxicity in Caco-2 cells (**Figure 3C**). Notably, addition of urea resulted in loss of conjugated BA-mediated protection of epithelial barrier integrity and cell viability (**Figure 3D-F**). We also exposed Caco-2 monolayers to combined BA pools in the presence or absence of urea and quantified the amount of unconjugated BAs that passed through the monolayer into the basolateral chamber by UPLC-MS. We observed significantly increased amounts of unconjugated BAs in the basolateral chamber in the presence of urea, providing further evidence that micelle formation prevents epithelial damage and permeability (**Figure 3G**). Together, our data provides evidence that unconjugated BAs lead to increased permeability across an intestinal epithelial monolayer by inducing cytotoxic effects. Further, these effects are prevented by the addition of conjugated BAs, which form BA micelles with unconjugated BAs in vitro (**Figure 3H**).

### BSH inhibition by AAA-10 prevents altered intestinal permeability and hepatic inflammation in CDAHFD-fed rats

Because conjugated BAs appear to protect against intestinal epithelial damage, we hypothesized that a decrease in microbial BA deconjugation by BSH might attenuate intestinal permeability in CDAHFD-fed rats. We recently reported the development of a covalent, gut-restricted, small molecule inhibitor of gut bacterial BSHs, AAA-10, a compound that effectively inhibits BSH activity and increases the abundance of conjugated BAs in vivo (**Figure 4A)** (Adhikari et al., 2021). We thus hypothesized that treatment of CDAHFD-fed rats with AAA-10 would prevent increased intestinal permeability. A dose of 10 mg/kg AAA-10 or vehicle control was administered via gavage twice a day for 7 days to CDAHFD-fed rats (**Figure 4B)**. AAA-10 administration led to ∼20-30 μM AAA-10 in cecal contents 48 hours and 1 week post-gavage (**Figure 4C**). No AAA-10 was detected in peripheral blood (**Figure S13A**), a finding that is consistent with our previous results (Adhikari et al., 2021) and indicates that this BSH inhibitor exhibits low systemic exposure. We found that AAA-10 reduced cecal BSH activity of CDAHFD-fed rats (**Figure 4D**) and significantly increased both the abundance of conjugated BAs as a group (**Figure 4E**) and the concentrations of individual conjugated BAs (**Figure S14**) by 1 week. Notably, we observed significant decreases in levels of portal venous LPS suggestive of increased intestinal barrier function in AAA-10-treated rats compared to vehicle-treated animals (**Figure 4F**). Furthermore, we observed normalization of ZO-1 localization and expression, suggesting that AAA-10 treatment prevented the development of intestinal permeability in CDAHFD-fed rats (**Figure 4G**).

**Figure 4.**
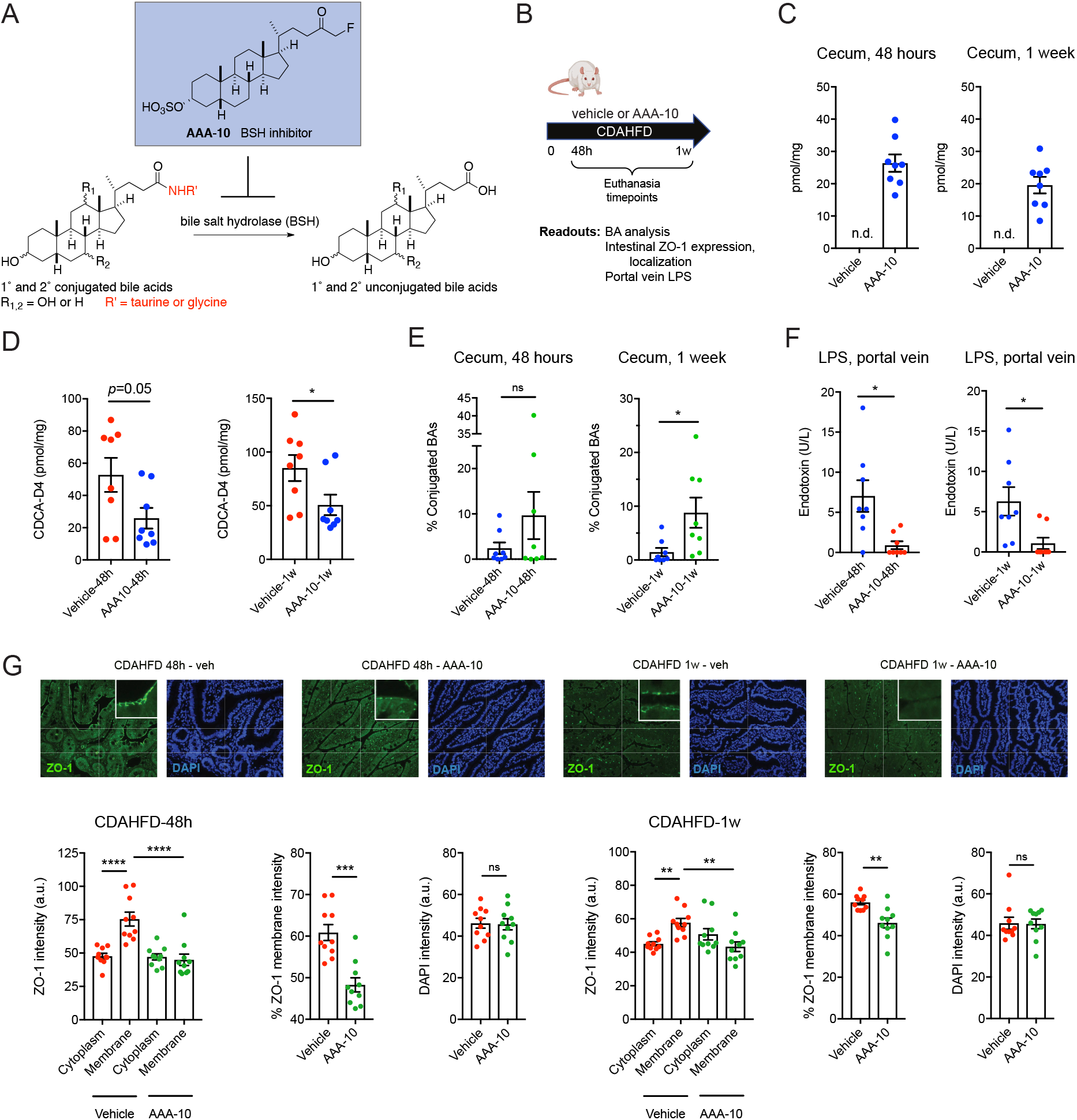
BSH inhibition increases intestinal conjugated BAs and prevents intestinal barrier dysfunction in CDAHFD-fed rats. (A) Schematic for the inhibition of bacterial BSH activity by the covalent pan-BSH inhibitor AAA-10 (Adhikari et al., 2021; Adhikari et al., 2020). (B) Schematic of rat intestinal permeability prevention experiment. Fats were fed CDAHFD and administered either AAA-10 (10 mg/kg/dose) or vehicle twice daily by gavage. Rats were euthanized at 48 hours or 1 week. Tissues and blood were collected for metabolite quantification and evaluation of intestinal injury markers (n=8 per for all timepoints). (C) UPLC-MS analysis of cecal contents of vehicle- and AAA-10-treated animals. Cecal AAA-10 levels were within the concentration range previously shown to be effective for inhibiting BSH activity (Adhikari et al., 2021; Adhikari et al., 2020). (D) Cecal BSH activity was reduced after 1 week of AAA-10 treatment. BSH activity was determined by quantifying conversion of GCDCA-d4 to CDCA-d4 in feces from vehicle- or AAA-10-treated animals (two-tailed Welch’s t-test). (E) Cecal conjugated BA abundance was significantly increased in AAA-10-treated rats after 1 week (two-tailed Welch’s t test, see Supplementary Information for concentrations of total and individual BAs in cecal contents at 1w). (F) Portal venous LPS levels were reduced in AAA-10 treated rats after 48h and 1w of AAA-10 treatment (two-tailed Welch’s t test). (G) AAA-10 treatment prevented aberrant ZO-1 subcellular localization. ZO-1 immunofluorescence and DAPI counterstaining of rat ileum from vehicle and AAA-10 treated CDAHFD-fed rats at 48h and 1w (n=10 intestinal cells quantified per group, see Methods for statistical analyses). Data not marked with asterisk(s) are not significant. \**p*<0.05, \*\**p*<0.005, \*\*\**p*<0.005, \*\*\*\**p*<0.0001. Bars represent mean ± SEM.

As intestinal permeability has been linked to translocation of intestinal products that further exacerbate hepatic inflammation (Chopyk & Grakoui, 2020), we evaluated the impact of 8 day AAA-10 treatment on liver phenotypes (**Figure 5A**). Consistent with our previous results, we found that the BA pool shifted toward conjugated BAs in AAA-10-treated compared to control-treated rats (**Figure S15**). Mild weight loss was observed in the AAA-10-treated animals (**Figure S16A**), consistent with our previous report that genetic removal of bacterial BSH causes altered metabolic phenotypes including reduced weight gain on a high fat diet (Yao et al., 2018). We did not observe significant weight loss in AAA-10-treated rats in the 7 day experiment (**Figure S13B**), and we observed similar changes in ZO-1 localization and expression in both experiments (**Figure 4G, Figure S16B**), suggesting that AAA-10-mediated protection against intestinal permeability occurred independent of weight loss. We also assessed the effect of AAA-10 or vehicle treatment on normal chow-fed rats and found no significant differences between the groups in liver or kidney function, body weight, or caloric intake (**Figure S17**), suggesting that AAA-10 itself was non-toxic and that differences between groups are directly related to changes in intestinal BA composition.

**Figure 5.**
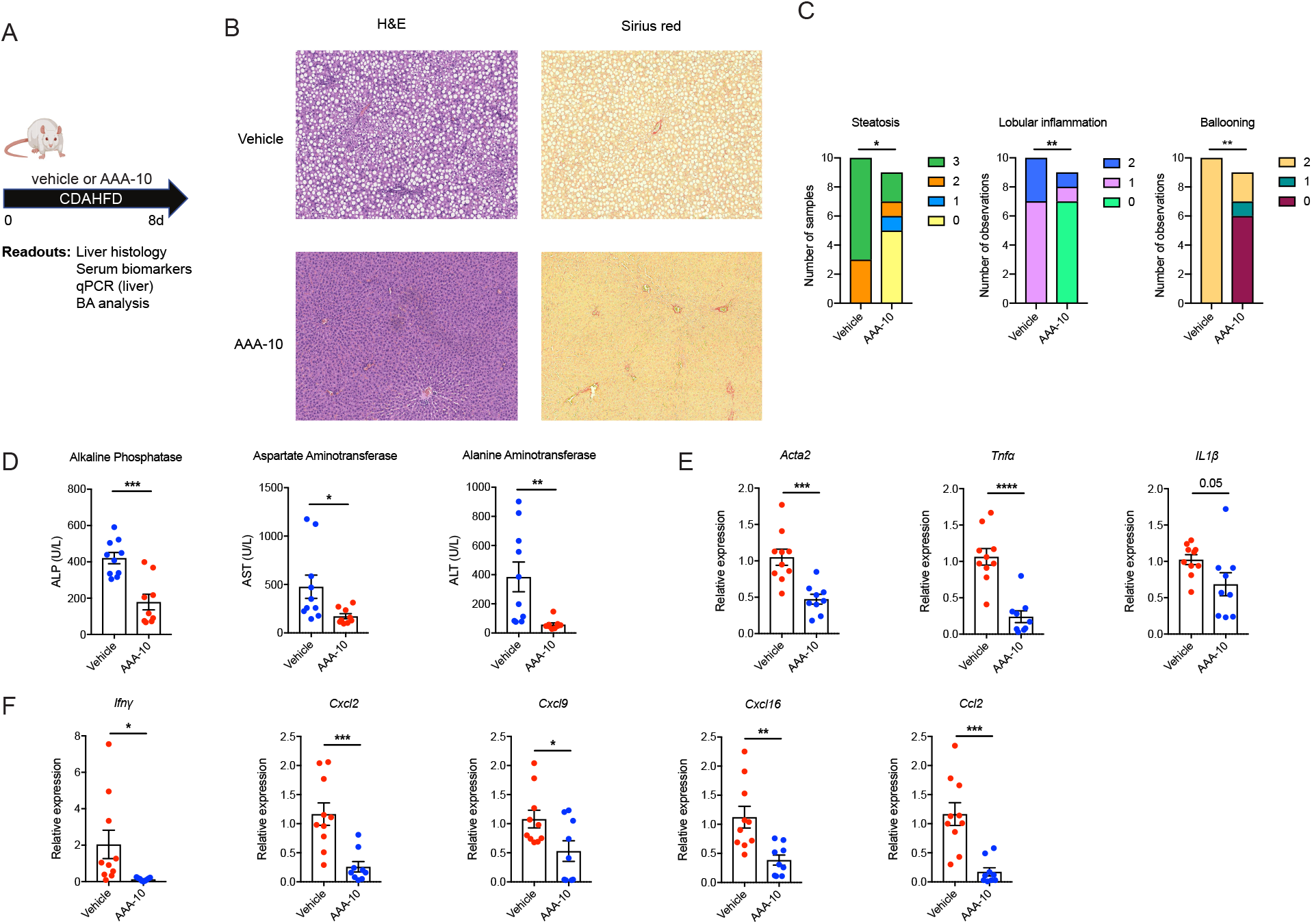
BSH inhibition prevents hepatic inflammation in CDAHFD-fed rats. Schematic of rat liver damage prevention experiment. Rats were fed CDAHFD and administered either AAA-10 (10 mg/kg/dose) or vehicle twice daily by gavage. Rats were euthanized after 8 days. Tissues and blood were collected for metabolite quantification and evaluation of liver injury markers (n=10 in vehicle group, n=9 in AAA-10 group for all timepoints). (B) AAA-10 treatment prevented development of hepatic steatosis and inflammation in CDAHFD-fed rats. Representative H&E staining of liver tissue is shown. (C) AAA-10 treatment prevented development of hepatic inflammation. Histologic scoring of steatosis, hepatocyte ballooning, and lobular inflammation in vehicle- and AAA-10-treated animals. Mann-Whitney test. (D) Serum ALT, AST, and ALP levels were decreased in AAA-10-treated compared to vehicle-treated rats, two-tailed Welch’s t test. (E, F) AAA-10 treatment attenuated hepatic gene expression of pro-inflammatory and pro-fibrotic genes in CDAHFD-fed rats as determined by RT-qPCR analysis of indicated genes. Two-tailed Welch’s t test. \**p*<0.05, \*\**p*<0.005, \*\*\**p*<0.0005, \*\*\*\**p*<0.0001. Bars represent mean ± SEM.

We observed substantial histologic improvements in AAA-10-treated rats compared to vehicle-treated controls. Specifically, AAA-10-treated rats exhibited significantly less hepatic steatosis, lobular inflammation, and hepatocyte ballooning (**Figure 5B,C**). Consistent with these results, we observed significant decreases in serum measures of hepatic inflammation including alanine transaminase (ALT), aspartate transaminase (AST), and alkaline phosphatase (ALP) in AAA-10-treated animals (**Figure 5D**). Finally, we observed decreased expression of pro-inflammatory and pro-fibrotic genes in treated rats (**Figure 5E-F**). Together, our findings demonstrate that treatment with the gut-restricted BSH inhibitor AAA-10 prevents the development of intestinal barrier dysfunction and liver damage in CDAHFD-fed rats.

### Glyco-conjugated BAs form protective micelles in vitro

While rodents possess only tauro-conjugated BAs, humans have a mixture of glyco- and tauro-conjugated BAs, with the former dominating the pool in a ratio of 3:1 (**Figure 6A**) (Ridlon et al., 2016). Further, microbial BSHs in the human intestine hydrolyze glyco-conjugated BAs to their unconjugated forms (Ridlon et al., 2016). In order to test whether glyco-conjugated BAs are also capable of forming micelles to sequester damaging unconjugated BAs, we determined critical micelle concentrations (CMCs) of these BA populations as described above (**Figure 3A, B**). We used an equimolar mix of (1) glyco-conjugated BAs (GCA, GDCA, GUDCA, GCDCA) or (2) combined BA pools containing the same unconjugated BAs as used above (βMCA, CA, DCA, UDCA, CDCA). Glyco-conjugated BAs exhibited a lower CMC than tauro-conjugated BAs (3.64 mM compared to 4.0 mM, respectively) (**Figure 6B, 3A**). Consistent with this observation and our previous results, the CMC of the combined pool with glyco-conjugated BAs was 3.60 mM, while the CMC of the combined pool of tauro-conjugated BAs was 4.2 mM (**Figure 6B, 3A**). We also performed direct visualization of micelles using negative staining-electron microscopy. At 5 mM concentration, no micelles were visible in the unconjugated BA pool while micelles were observed in the glyco-conjugated and combined BA pools, consistent with our CMC determinations (**Figure 6C**). Together, these results suggest that glyco-conjugated BAs form micelles with unconjugated BA species with equal to or greater efficiency than tauro-conjugated BAs.

**Figure 6.**
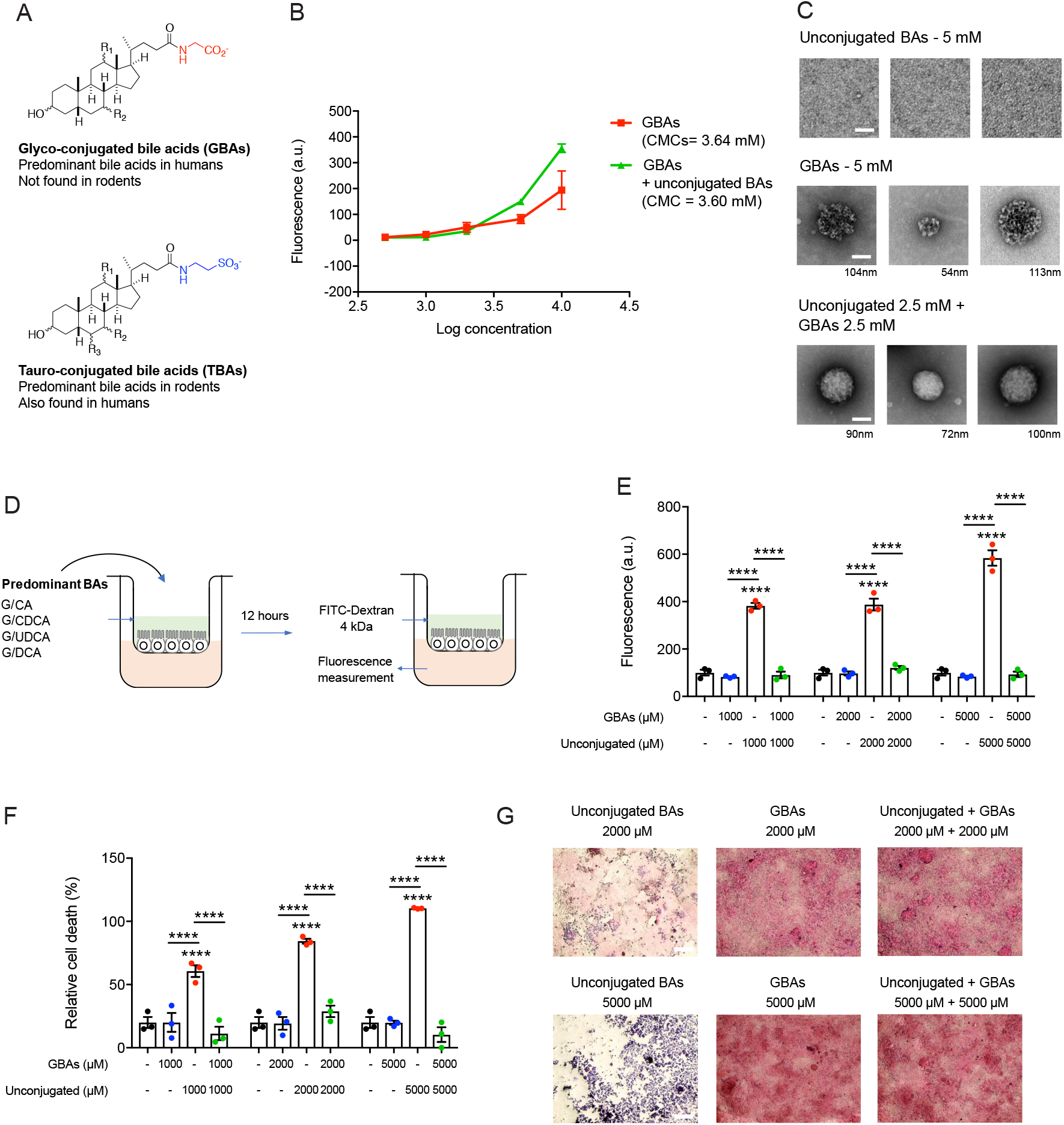
Predominant human glyco-conjugated BAs sequester unconjugated BAs in micelles and prevent epithelial damage in vitro. (A) Structures of glyco- and tauro-conjugated BAs (GBAs and TBAs, respectively). While TBAs are found in both rodents and humans, GBAs are the predominant conjugated BAs in humans and are not present in rodents. (B) Pools of GBAs alone (equimolar concentrations of GCA, GCDCA, GUDCA, GDCA) and GBAs plus unconjugated BAs (equimolar concentrations of glyco-conjugated BAs with βMCA, CA, CDCA, UDCA, DCA) exhibited CMCs of 3.64 mM and 3.60 mM, respectively, indicating efficient micelle formation (n=3 per group). (C) EM images of micelles formed from BA pools at indicated concentrations. Similar to TBAs (see Figure 3), while detectable micelles were visible in the GBAs alone or unconjugated BAs plus GBA pools, no detectable micelles were visible in the unconjugated BAs pool. Scale bar=50 nm, diameter of micelle indicated beneath each image. (D) Schematic of in vitro permeability experiment. Equimolar concentrations of GBAs, unconjugated BAs, and combined BA pools were added to the apical chamber of transwell system containing Caco-2 monolayer and incubated for 12 hours before addition of FITC-Dextran 4kDa and fluorescence measurement from basolateral chamber. (E) GBAs protected against unconjugated BA-induced Caco-2 monolayer permeability at physiologic concentrations (n=3 per group, see Methods for statistical analyses). (F) GBAs protected epithelial monolayers from unconjugated BA-induced cell death. Cell viability of Caco-2 cells measured by MTT assay (n=3 per group, see Methods for statistical analyses). (G) GBAs protected the physical integrity of epithelial monolayers from unconjugated BA-induced damage. Representative light microscopy images of H&E-stained Caco-2 monolayers after exposure to BA pools. Scale bar=20 μm. Unless otherwise specified, all experiments were performed in triplicate. See Methods for statistical analyses. Data not marked with asterisk(s) are not significant \*\*\*\**p*<0.0001. Bars represent mean ± SEM.

Next, we sought to determine whether glyco-conjugated BAs would also protect the intestinal epithelium from the damaging effects of unconjugated BAs. Using transwell-differentiated Caco-2 cells, we found that unconjugated BA-induced epithelial monolayer damage was prevented by addition of an equimolar concentration of glyco-conjugated BAs (**Figure 6D, E**). Glyco-conjugated BAs alone did not disrupt monolayer integrity at any concentrations tested (**Figure 6E**). Similarly, while unconjugated BAs were toxic to cells, equimolar addition of glyco-conjugated BAs abrogated this effect (**Figure 6F**). Visualization of Caco-2 monolayers by hematoxylin and eosin staining further confirmed that the toxic effects of unconjugated BAs were largely prevented by addition of glyco-conjugated BAs (**Figure 6G**). Together, these results suggest that the predominant human-derived glyco-conjugated BAs are also capable of sequestering unconjugated BAs in micelles and protecting against epithelial permeability.

## Discussion

In this study, we show that the onset of intestinal permeability is associated with decreased intestinal and portal conjugated BA abundance in a diet-induced animal model of intestinal barrier and liver damage. Furthermore, we demonstrate a mechanism by which conjugated BAs protect intestinal epithelial barriers from damage in vitro via the sequestration of unconjugated BAs in micelles. We then demonstrate that inhibition of gut bacterial BSH activity increases the abundance of cecal conjugated BAs and prevents the development of intestinal barrier dysfunction and liver inflammation in vivo. Finally, we show that both tauro-conjugated BAs and glyco-conjugated BAs protect against unconjugated BA-induced epithelial barrier damage in vitro. Because glyco- and tauro-conjugated BAs are the two conjugated BA forms found in humans, our data suggest that conjugated BAs could be protective in the human gut.

The CDAHFD-fed rat model we utilized here to investigate pathogenic intestinal permeability is a rodent model of chronic liver disease, including NAFLD and NASH progression (Longo et al., 2020). Gut dysbiosis is a central feature of human NAFLD/NASH and has been proposed to contribute to the development of these conditions (Boursier et al., 2016; Loomba et al., 2019). Likewise, changes in serum BA profiles have been reported to correlate with disease severity in NAFLD/NASH patients (Mouzaki et al., 2016; Nimer et al., 2020; Puri et al., 2018). However, causal connections between gut bacteria, bile acid metabolism, and NAFLD/NASH in humans have not yet been established. Our data raise the possibility that bacterial BA deconjugation may be causally contributing to NAFLD/NASH progression. Indeed, we observed higher levels of BSH activity in CDAHFD-treated rats compared to control animals. Future analyses will help uncover whether correlations exist between fecal BA levels, BSH activity, and disease severity in human patients. In addition, further studies in different animal models of pathogenic intestinal permeability will reveal whether pharmacological BSH inhibition can prevent or ameliorate gut barrier damage. Nonetheless, the finding that BSH inhibition prevents intestinal permeability and liver damage in choline-deficient high fat diet-fed rodents, a clinically relevant a diet-induced model of liver cirrhosis, suggests that modulating the in vivo BA pool by selective targeting of the gut microbiota could be investigated as a treatment path for diseases characterized by intestinal barrier disruption.

While intestinal barrier function has been implicated in the pathogenesis of a variety of diseases, the molecular mechanisms that trigger or protect against intestinal permeability defects are incompletely defined (Gupta et al., 2020). It has been shown that hydrophilic BAs, including conjugated BAs, protect against cytotoxicity induced by hydrophobic bile acids, including many unconjugated species (Araki et al., 2003; Di Ciaula et al., 2017; Hegyi, Maleth, Walters, Hofmann, & Keely, 2018). However, prior to our work, the connection between the efficiency of micelle formation by conjugated BAs and their ability to protect against unconjugated BA-induced epithelial barrier damage had not been established. Here, we demonstrate that conjugated BAs form micelles with unconjugated BAs at physiologically relevant concentrations in vitro. Micelle formation leads to sequestration of hydrophobic, unconjugated BAs away from epithelial cells and prevents cell death and tight junction dysfunction. These data demonstrate that distinct classes of BAs elicit differential effects on intestinal epithelial permeability and delineate a mechanism by which one of these classes, conjugated BAs, protects against epithelial damage in vitro. Whether this mechanism is operable in vivo, however, is not yet known. Cholesterol, phospholipids, and lipid hydrolysis products form mixed micelles with bile acids in bile and in the intestine (Hofmann, 1999). The presence of these other compounds could alter micelle formation and thus the protective effects of conjugated BAs observed in vitro. To our knowledge, there are currently no selective means of preventing micelle formation in vivo. Nonetheless, by inhibiting BSH activity using AAA-10, we selectively shifted the in vivo BA pool toward conjugated BAs. These experiments showed that, consistent with our in vitro data, conjugated BAs protect against epithelial barrier damage in vivo.

The treatment of CDAHFD-fed rats with a gut-restricted BSH inhibitor not only prevented intestinal permeability but also protected against the development of hepatic inflammation and steatosis. The prevention of liver damage is likely multifactorial, and effects including decreased translocation of bacterial products such as LPS as well as mild weight loss may contribute. The weight loss observed is likely a result of BSH-dependent metabolic changes as we have previously reported (Yao et al., 2018). Interestingly, germ-free mice, which lack BSH and therefore have high levels of intestinal conjugated BAs, are resistant to development of hepatic steatosis (Brandl & Schnabl, 2017). Moreover, colonization of germ-free mice on a HFD with BSH-containing bacteria induces hepatic steatosis, a phenotype that can be prevented by deletion of BSH (Yao et al., 2018). Our results here demonstrate that shifting the in vivo BA pool to enrich for conjugated BAs can not only prevent pathogenic intestinal permeability but can also prevent fat deposition and inflammation in the host liver.

Overall, our findings link changes in the intestinal BA pool, specifically conjugated BAs, with maintenance of the intestinal epithelial barrier in vivo. Our study also reveals that bacterial BSH enzymes regulate metabolites that control gut barrier integrity, thereby providing a potential mechanistic link between the microbiome and the development of pathogenic intestinal permeability. Our findings suggest that strategies that reduce BSH activity in the human gut microbiome could be developed as novel paradigms to treat intestinal barrier dysfunction.

## Materials and Methods

### Animals

8-week old Wistar rats were purchased from Charles River Laboratories (Wilmington, MA) and housed in a specific pathogen-free environment (maximum four per cage). After 10 days of acclimation, rats were initiated on either a control high-fat diet (60 kcal% fat; Research Diets D12492) or CDAHFD (L-amino acid diet with 60 kcal% fat with 0.1% methionine without added choline; Research Diets A06071302) *ad libitum* for either 48 hours or 1 week (either 7 or 8 days). At time of sacrifice, rats were anesthetized using 100 mg/kg of ketamine and 10 mg/kg of xylazine intraperitoneally followed by portal vein blood draw and terminal cardiac puncture.

### In vivo intestinal permeability assay

Intestinal permeability was assessed by in vivo fluorescein isothiocyanate (FITC)-dextran (FD4; Sigma-Aldrich, St. Louis, MO) permeability assay as described previously (Woting & Blaut, 2018). Rats were fasted for 4 hours and then had blood collected via tail vein puncture to assess background fluorescence. Rats were then gavaged with 0.4 mg/g body weight FITC-dextran 4kDa solution and blood was collected by terminal cardiac puncture. Fluorescence intensity in the serum determined at 530 nm with excitation at 485 nm. Relative fluorescence units determined by subtracting the PBS blank fluorescence from all samples and then subtracting the pre-gavage fluorescence from the post-gavage fluorescence. FITC-dextran concentrations were determined from a standard curve generated by serial dilutions of FITC-dextran.

### AAA-10 treatment

After initiation of CDAHFD diet, rats were split into two groups and were gavaged twice a day with either 10 mg/kg of AAA-10 dissolved in 5% Captisol (Ligand, San Diego, CA) and 5% DMSO in PBS or an equal volume of 5% Captisol and 5% DMSO in PBS.

### RNA extraction and RT-qPCR

Total RNA was extracted from liver tissue using TRizol (Invitrogen) according to the manufacturer’s instructions and subsequently treated with DNase I (Promega). cDNA was generated using the RevertAid First Strand cDNA Synthesis kit according to manufacturer’s instructions (Thermo Fisher) and RT-qPCR was performed using the Power SYBR Green Master Mix Kit (Thermo Fisher). Expression of *GAPDH* was used to standardize the samples, and the results are expressed as a ratio relative to control. The genes for which expression was determined and the primer sequences used in this study can be found in **Supplementary Table 1**.

### Serum processing

A cardiac terminal blood withdrawal was performed at the time of sacrifice. Blood was allowed to clot for 2 hours at room temperature before centrifugation at 2,000 rpm for 10 minutes at 4°C. Serum was isolated and stored at -80°C. Biochemical markers of liver injury were measured, including alkaline phosphatase, alanine aminotransferase, and aspartate aminotransferase (DRI-CHEM 4000 Analyzer, Heska).

### Histology and immunofluorescence

Formalin fixed paraffin embedded (FFPE) liver and ileum tissue were sectioned at a thickness of 4 um. Slides were stained with hematoxylin and eosin at the MGH Cytopathology Core. Slides were stained with Sirius Red for fibrosis staging. Slides were blindly reviewed by a blinded pathologist to grade steatosis, inflammation, and fibrosis, using criteria modeled on human NASH histologic scoring systems. Similarly, for intestine, slides were evaluated for intestinal inflammation, epithelial hyperplasia, and goblet cell loss. For immunofluorescence, sections were stained for ZO-1 (Abcam, Cambridge, MA) with detection by appropriate secondary antibodies labeled with Alexa488 according to the manufacturer’s instructions. Hydroxyproline was quantified by high-performance liquid chromatography analysis as previously described (Hutson, Crawford, & Sorkness, 2003).

For ZO-1 staining intensity measurement, images were processed and analyzed with Adobe Photoshop CC software. Matched images were taken with the same exposure and were processed and analyzed identically. The two-dimensional image intensities of ZO-1 fluorescence (in pixels) of n=10 randomly selected cells in micrographs from each condition were measured using Photoshop software (Adobe) and plotted in GraphPad Prism.

### LPS measurement assay

Bacterial endotoxin measurement in portal serum samples was performed using the Lonza Pyrogent turbidometric LAL assay kit (Lonza, N383) according to manufacturer’s instructions. Briefly, serial dilutions of prepared serum samples were made using the LAL reagent water provided in the kit, followed by incubation for 15 min at 37ºC. Samples were then mixed with the reconstituted Pyrogent reagent from the kit, followed by kinetic turbidometric analyses in a SpectraMax M5 plate reader (Molecular Devices, San Jose, CA) at the ICCB-Longwood Screening Facility at HMS. LPS amount in samples was deduced using standard curves generated from stock provided in the kit and linear regression calculation.

### Bile acid analysis

BA analyses were performed using a previously reported method (Yao et al., 2018).

#### Reagents

Stock solutions of all bile acids were prepared by dissolving compounds in molecular biology grade DMSO (VWR International). These solutions were used to establish standard curves. HPLC grade solvents were used for preparing and running UPLC-MS samples.

#### Extraction

Rat cecal and human fecal samples (approximately 50 mg each) were collected in pre-weighed lysis tubes. The lysis tubes contained ceramic beads to allow for homogenization (Precellys lysing kit tough micro-organism lysing VK05 tubes for rat cecal and human fecal samples; Bertin technologies, Montigny-le-Bretonneux, France). 400 μL of methanol (MeOH) was added to rat cecal and human feces and the tubes were homogenized in a MagNA Lyser (6000 speed for 90 s*2, 7000 speed for 60 s). For portal venous samples, serum (100 μL) was collected in eppendorf tubes, followed by addition of 100 μL MeOH. Samples were vortexed and frozen at -20°C until further analysis. Cell culture media was diluted 1:1 in MeOH. All MeOH-extracted samples were centrifuged at 4°C for 30 min at 15,000 rpm. The supernatant was diluted 1:1 in 50% MeOH/water and centrifuged again at 4°C for 30 min at 15000 rpm. The supernatant was transferred into mass spec vials and injected into the UPLC-MS. Total bile acids were then calculated by adding all detected and measured bile acids. The limits of detection for individual bile acids have been described previously (Chaudhari, Harris, et al., 2021; Chaudhari, Luo, et al., 2021; Yao et al., 2018).

### Caco-2 cell culture and differentiation

Caco-2 cells were obtained from American Type Culture Collection (Manassas, VA). Cells were maintained in Minimum Essential Medium (MEM) with GlutaMAX and Earle’s Salts (Gibco, Life Technologies, UK). Cell culture media were supplemented with 10% fetal bovine serum (FBS), 100 units/mL penicillin, and 100 μg/mL streptomycin (GenClone). Cells were grown in FBS- and antibiotic-supplemented ‘complete’ media at 37°C in an atmosphere of 5% CO_2_. Undifferentiated Caco-2 cells were seeded in 24-well plate transwells (0.4 μM pore size, Costar) at 200,000 cells per transwell. Media was changed on days 4, 8, 12, 16, and 18 to differentiate Caco-2 cells in vitro (Lea, 2015). On day 21, fully differentiated and polarized cells were used for permeability and toxicity assays.

### H&E staining and light microscopy of transwells

Differentiated Caco-2 cells in transwells were stained with H&E stain (Abcam, ab245880) according to manufacturer’s instructions. Briefly, transwells were washed with PBS, followed by staining with hematoxylin for 5 min. Transwells ere rinsed in twice in dH_2_O to remove excess stain, followed by addition of bluing reagent and incubation for 10-15 sec. Two rinses in dH_2_O was performed prior to addition of ethanol (Sigma) to remove excess reagent. Ethanol was removed, and adequate Eosin was added for 2-3 min for the counter stain. Transwells were dehydrated in ethanol 2-3 times, dried, and imaged using an Evos XL Core microscope (Invitrogen) attached to camera at the ICCB-Longwood Screening Facility at HMS. Cells were imaged from the top, and all images were taken at the same magnification and light intensity. For each treatment, n=2 transwells were imaged, providing 4 images per transwell. Representative images are shown in **Figures 2, 3**, and **6**.

### In vitro bile acid treatments

Caco-2 cells (undifferentiated in 96-well plates or day 21 to 25 of differentiation in transwells) were treated with bile acid mixtures for 12-16 hours prior to assays. Diluted stocks of bile acid standards or undiluted methanol-extracted cecal contents were added in complete media. Transcytosis of bile acids was measured by drying basolateral media in a speed vac followed by resuspending media in 1:1 methanol/water, transferred into mass spectrometry vials and injected onto the ultra-high performance liquid chromatography-mass spectrometer (UPLC-MS).

### Caco-2 permeability assay

Epithelial integrity by FITC-dextran permeability assay was performed as described previously (Chaudhari, Harris, et al., 2021). Briefly, differentiated Caco-2 epithelial integrity was assayed by measuring passive diffusion of 4 kDa FITC-Dextran (Sigma Aldrich) added at a concentration of 5 μM to the apical chamber in 100 μL PBS, while the basolateral chamber contained 500 μL PBS. Diffusion from the apical to basolateral side was measured by fluorescence reading in PBS on the basolateral side of the transwell system using a SpectraMax M5 plate reader (Molecular Devices, San Jose, CA) at the ICCB-Longwood Screening Facility. Fluorescence reading was normalized to the control.

### Caco-2 cell viability assay

Caco-2 (undifferentiated in 96-well plates or day 21 to 25 of differentiation in transwells) cell viability was measured using the MTT assay (Abcam, ab211091) according to manufacturer’s instructions. Cells were treated with compounds for 12-16 hours prior to viability determination. Briefly, cell culture media was replaced with the MTT reagent and incubated for 3 hours at 37ºC. Following incubation, the MTT reagent was replaced with the MTT solvent and incubated for 15 min, followed by analysis in a SpectraMax M5 plate reader (Molecular Devices, San Jose, CA) at the ICCB-Longwood Screening Facility at HMS. Absorbance reading at 690 nm, as a measure of viability, was normalized to the control.

### Critical micelle concentration (CMC) assay

CMC determination of groups of bile acids was performed using a previously described assay using coumarin 6 as a fluorescent probe with minor adaptations (Fluksman & Benny, 2019). Briefly, 6 mM coumarin 6 (Sigma) in dichloromethane (Sigma) was added to Eppendorf tubes and allowed to evaporate for 30 min in a chemical hood. 400 μL of equimolar mixtures of bile acids at various concentrations to be tested were added to tubes and rotated overnight at room temperature in the dark (unconjugated BAs: beta-muricholic acid [βMCA], cholic acid [CA], deoxycholic acid [DCA], ursodeoxycholic acid [UDCA], chenodeoxycholic acid [CDCA]); tauro-conjugated BAs: tauro-beta-muricholic acid [TβMCA], tauro-cholic acid [TCA], tauro-deoxycholic acid [TDCA], tauro-ursodeoxycholic acid [TUDCA], tauro-chenodeoxycholic acid [TCDCA]); glyco-conjugated BAs: glyco-cholic acid [GCA], glyco-deoxycholic acid [GDCA], glyco-ursodeoxycholic acid [GUDCA], glyco-chenodeoxycholic acid [GCDCA]). The next day, 200 μL of the solution was transferred to black 96 well plates (Costar), and fluorescence intensity at 480/530 was measured using a SpectraMax M5 plate reader (Molecular Devices, San Jose, CA) at the ICCB-Longwood Screening Facility. Fluorescence intensity was plotted against the logarithm of the corresponding concentration and the CMC was determined by the intersection of the two tangents created in the graph.

### Electron microscopy

Caco-2 cells seeded on transwells were washed in PBS prior to fixation in FGOP (Formaldehyde-Glutaraldehyde-Picric acid) fixative. Samples were fixed overnight in a mixture of 1.25% formaldehyde, 2.5 % glutaraldehyde, and 0.03% picric acid in 0.1 M sodium cacodylate buffer, pH 7.4. FGP fixative was diluted in PBS 1:1 before applying to the apical and basolateral chamber of transwells. Fixed samples were stored at 4°C until further processing and imaging by the Electron Microscopy Core at Harvard Medical School. All electron microscopy imaging was performed by a blinded investigator.

For transmission electron microscopy, fixed tissues were washed with 0.1M sodium cacodylate buffer and post fixed with 1% osmium tetroxide/1.5% potassium ferrocyanide (in H_2_O) for 2 hours. Samples were then washed in a maleate buffer and post fixed in 1% uranyl acetate in maleate buffer for 1 hour. Tissues were then rinsed in ddH_2_O and dehydrated through a series of ethanol (50%, 70%, 95%, (2x)100%) for 15 minutes per solution. Dehydrated tissues were put in propylene oxide for 5 minutes before they were infiltrated in epon mixed 1:1 with propylene oxide overnight at 4C. Samples were polymerized in a 60ºC oven in epon resin for 48 hours. They were then sectioned into 80nm thin sections and imaged on a JEOL 1200EX Transmission Electron Microscope.

For negative staining for visualization of micelles, the sample was diluted in water and adsorbed onto a glow-discharged carbon grid. Once the specimen was adsorbed on to the film surface, the excess liquid was blotted off using a filter paper (Whatman #1) and the grid was floated on a small drop (∼5 μl) of staining solution (0.75% uranyl acetate). After 20 seconds the excess stain was blotted off and the sample was air dried briefly before it was examined in the TEM.

### BSH activity assay

BSH activity was quantified using a modified version of a previously described method (Adhikari et al., 2020). Briefly, fresh cecal contents (approximately 20 mg) were diluted in PBS to obtain a concentration of 1 mg/mL 100 μM glycochenodeoxycholic acid-d4 (GCDCA-d4) was added to the mixture and incubated at 37ºC for 30 minutes, then frozen in dry ice for 5 minutes and stored at -80ºC until further analysis. On thawing, the mixture was diluted with an equal volume of methanol and the slurry was centrifuged at 12,500 x g for 10 minutes. The supernatant was removed into a clean Eppendorf tube and centrifuged again. The supernatant was transferred to mass spectrometry vials and samples were analyzed as described in Bile Acid Analysis in Supplementary Information.

### Statistical Analyses

Data was quantified using software linked to indicated instruments and plotted in GraphPad Prism 7. Statistical analyses were performed using GraphPad Prism and Microsoft Excel software. Statistical significance was assessed using Student’s or Welch’s t tests, one-way or two-way ANOVAs followed by multiple comparisons tests, and Mann-Whitney tests wherever appropriate. **Figures 1E, F, 4G**, and **S16**: Intestines from all animals were stained, n=10 intestinal cell images per group were analyzed. For ZO-1 intensity, one-way ANOVA followed by Tukey’s multiple comparison test, for %ZO-1 membrane intensity and DAPI intensity, two-tailed Welch’s t test. Figures 2A, B: one-way ANOVA followed by Dunnett’s multiple comparison test for treatment vs. DMSO, one-way ANOVA followed by Tukey’s multiple comparison test for comparing treatments. **Figures 2D, E, 3D, E, G, 6E, F**: 2-way ANOVA followed by Sidak’s multiple comparisons test.

### Ethics

Animals received humane care per criteria outlined in the Guide for the Care and Use of Laboratory Animals by the National Academy of Sciences (National Institutes of Health publication 86-23, revised 1985) and in accordance with the Massachusetts General Hospital Institutional Animal Care and Use Committee guidelines (Protocol 2007N000113).

## Author Contributions

D.K.L., S.N.C., R.T.C, and A.S.D. conceived the project and designed the experiments. D.K.L., M.S., Y.L, and S.C.B. performed all animal experiments, immunofluorescence, staining, and transcriptional analyses on rat tissues. S.N.C. performed the cell culture experiments, BA profiling, and preparation for electron microscopy. S.S. and L.Z. performed pathology scoring of rat tissue. D.K.L., S.N.C., and A.S.D. wrote the manuscript. All authors edited and contributed to the critical review of the manuscript.

## Acknowledgements

We are indebted to members of the Devlin, Chung, and Tanabe groups for helpful discussions. We thank the HMS Electron Microscopy core for technical support and advice. We are grateful to the human patients who participated in this study. The research was supported by National Institutes of Health (NIH) grant R35 GM128618 (A.S.D.), an Innovation Award from the Center for Microbiome Informatics and Therapeutics at MIT (A.S.D), a grant from Harvard Digestive Diseases Center (supported by NIH grant 5P30DK034854-32) (A.S.D), a John and Virginia Kaneb Fellowship (A.S.D), a Quadrangle Fund for the Advancement and Seeding of Translational Research at Harvard Medical School (Q-FASTR) grant (A.S.D), an HMS Dean’s Innovation Grant in the Basic and Social Sciences (A.S.D), and the MGH Research Scholars Program (R.T.C.). D.K.L. was supported by National Institutes of Health T32 training grant (5T32DK007191). S.N.C. acknowledges an American Heart Association Postdoctoral Fellowship. Work by Z.Z. and S.H.S. was funded in part by the Harvard Medical School Foundry. The rat images in the figures were provided by Biorender.

## Competing Interests Statement

The authors declare the following competing financial interest(s): A.S.D. is an *ad hoc* consultant for Takeda Pharmaceuticals and Axial Therapeutics. The other authors declare that no competing interests exist.

## Data Transparency Statement

All data generated or analyzed during this study are included in this article and its Supplementary Information.

## Notes

### Summary of Updates

Revised version of manuscript.

