## Supplementary Information for "Inhibition of microbial deconjugation of micellar bile acids protects against intestinal permeability and liver injury"

Figure S1

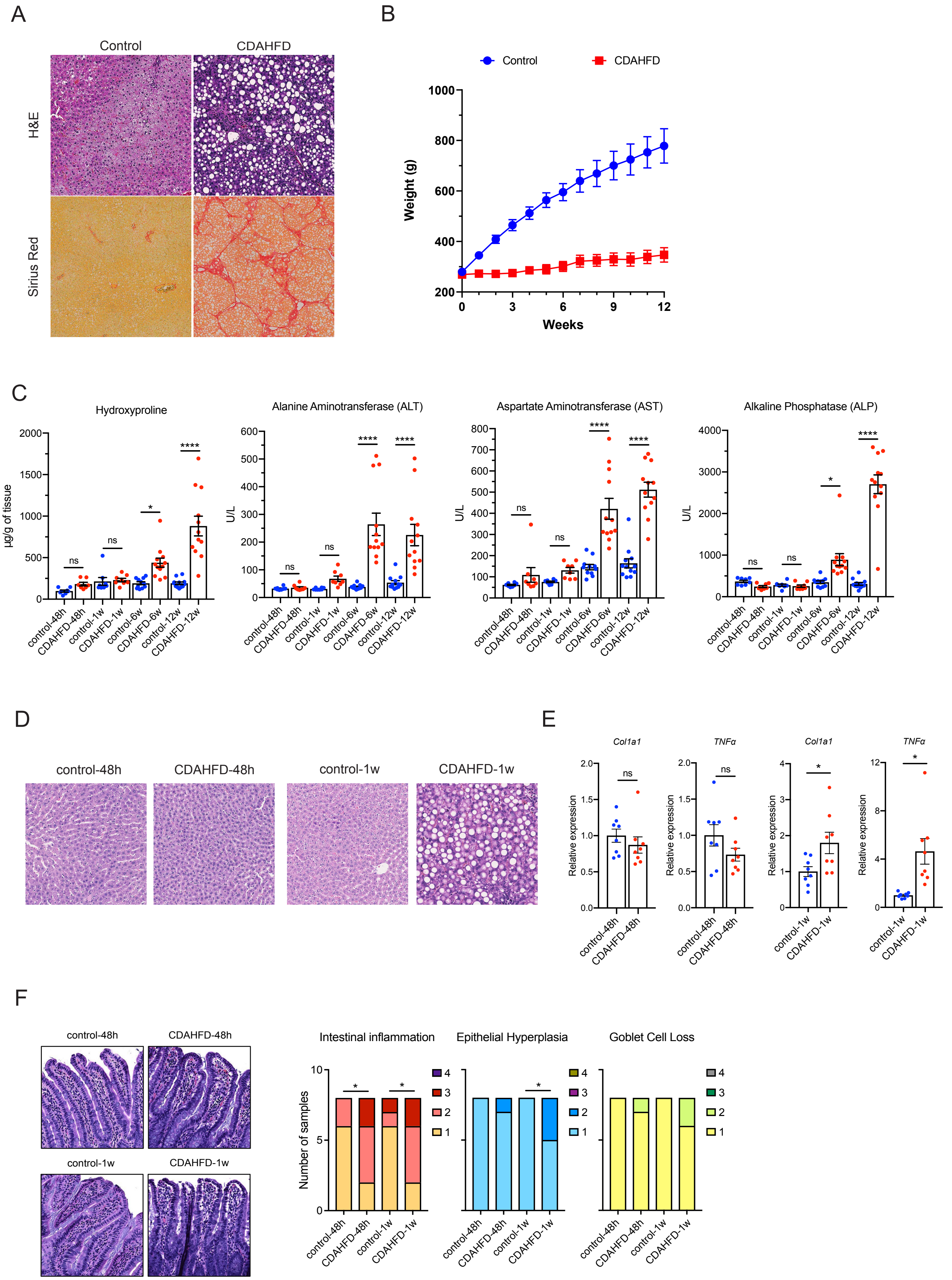
A
Control
CDAHFD
H&E
Sirius Red
B
Control
CDAHFD
Weight (g)
Weeks
C
Hydroxyproline
µg/g of tissue
Alanine Aminotransferase (ALT)
U/L
Aspartate Aminotransferase (AST)
U/L
Alkaline Phosphatase (ALP)
U/L
control-48h
CDAHFD-48h
control-1w
CDAHFD-1w
control-6w
CDAHFD-6w
control-12w
CDAHFD-12w
D
control-48h
CDAHFD-48h
control-1w
CDAHFD-1w
E
Col1a1
TNFα
Col1a1
TNFα
Relative expression
F
control-48h
CDAHFD-48h
control-1w
CDAHFD-1w
Intestinal inflammation
Epithelial Hyperplasia
Goblet Cell Loss
Number of samples

**Supplementary Figure 1. Temporal development of liver damage in CDAHFD-fed rats**. (A) CDAHFD-fed rats developed cirrhosis after 12 weeks of diet. Representative hemotoxylin and eosin (H&E) and Sirius Red staining of livers from CDAHFD-fed and control rats after 12 weeks of diet. (B) Body weight of CDAHFD-fed and control rats, measured weekly. (C) CDAHFD-fed rats developed progressive hepatic inflammation and fibrosis as demonstrated by increased ALT, AST, alkaline phosphatase, and hepatic hydroxyproline from livers of control and CDAHFD-fed rats. n=8 per group for 48h and 1w timepoints (n=12 per group for 6w and 12w timepoints, one-way ANOVA followed by Tukey's multiple comparison test). (D) Histologic evidence of hepatic inflammation was evident after 1 week of CDAHFD but not after 48h. Representative H&E staining of liver tissue from control and CDAHFD-fed rats at indicated timepoints. (E) Expression of fibrosis and pro-inflammatory genes (hepatic *Col1a1* and *TNFα*, respectively) were increased after 1 week of CDAHFD but not after 48h as determined by qPCR (n=8 per group, two-tailed Welch's t test). (F) CDAHFD diet induced intestinal inflammation. Representative hematoxylin and eosin (H&E) staining of ileum from control and CDAHFD-fed rats with pathology scores (n=8 per group, Mann-Whitney test). ns = not significant, *$p$<0.05 **$p$<0.005, ***$p$<0.001, ****$p$<0.0001. Bars represent mean ± SEM.

Figure S2

Rat cecal BAs, 48h

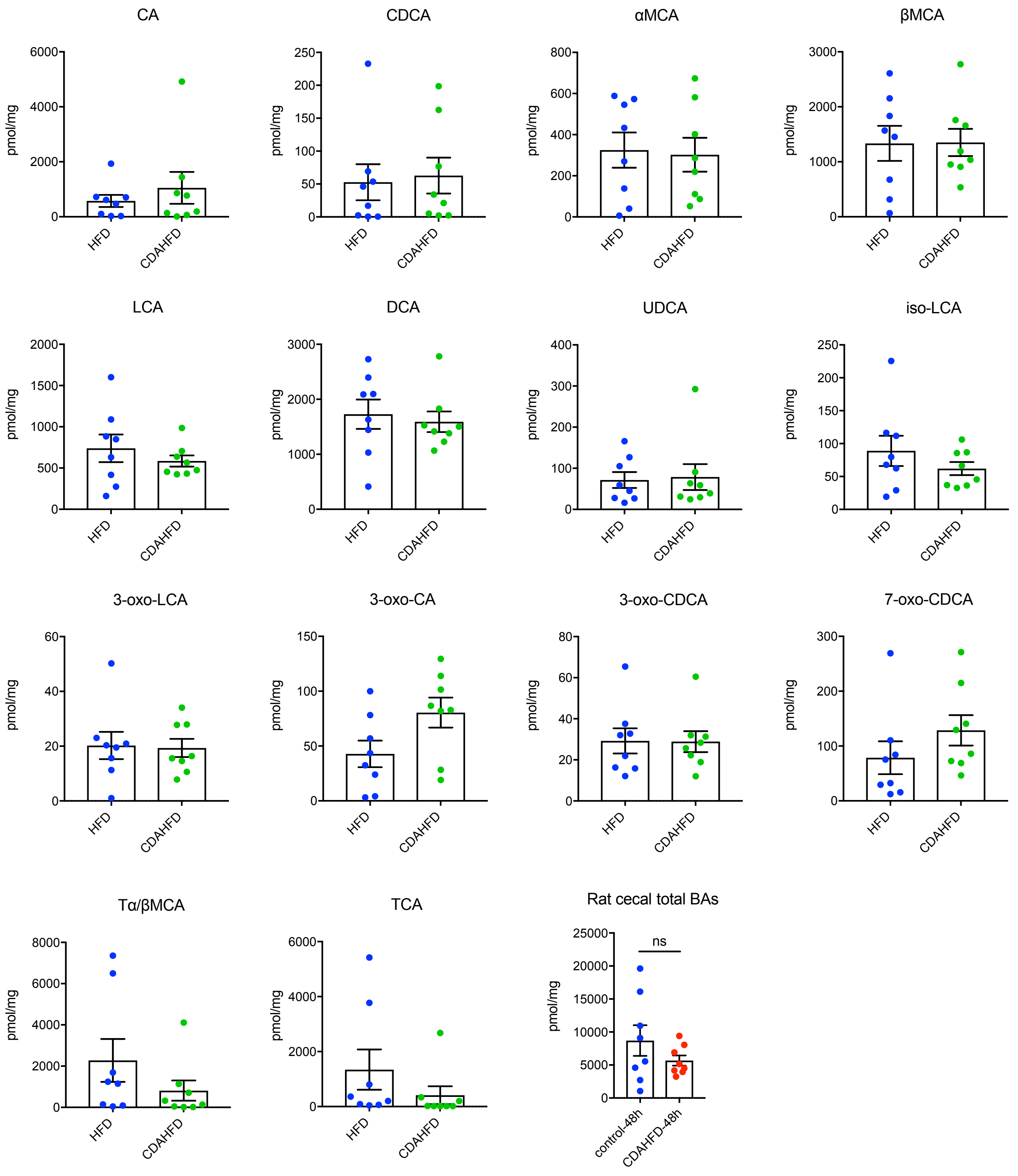

**Supplementary Figure 2**. **Bile acid concentrations in cecal contents of rats 48h post HFD control or CDAHFD diet intervention**. Bile acids were quantified using UPLC-MS. All bile acids with measurable concentrations above the limit of detection are shown (n=8 per group, two-tailed Welch's t test, data not marked with asterisk(s) were not significant). ns = not significant, Bars represent mean ± SEM.

### Figure S3

#### Rat cecal BAs, 1w

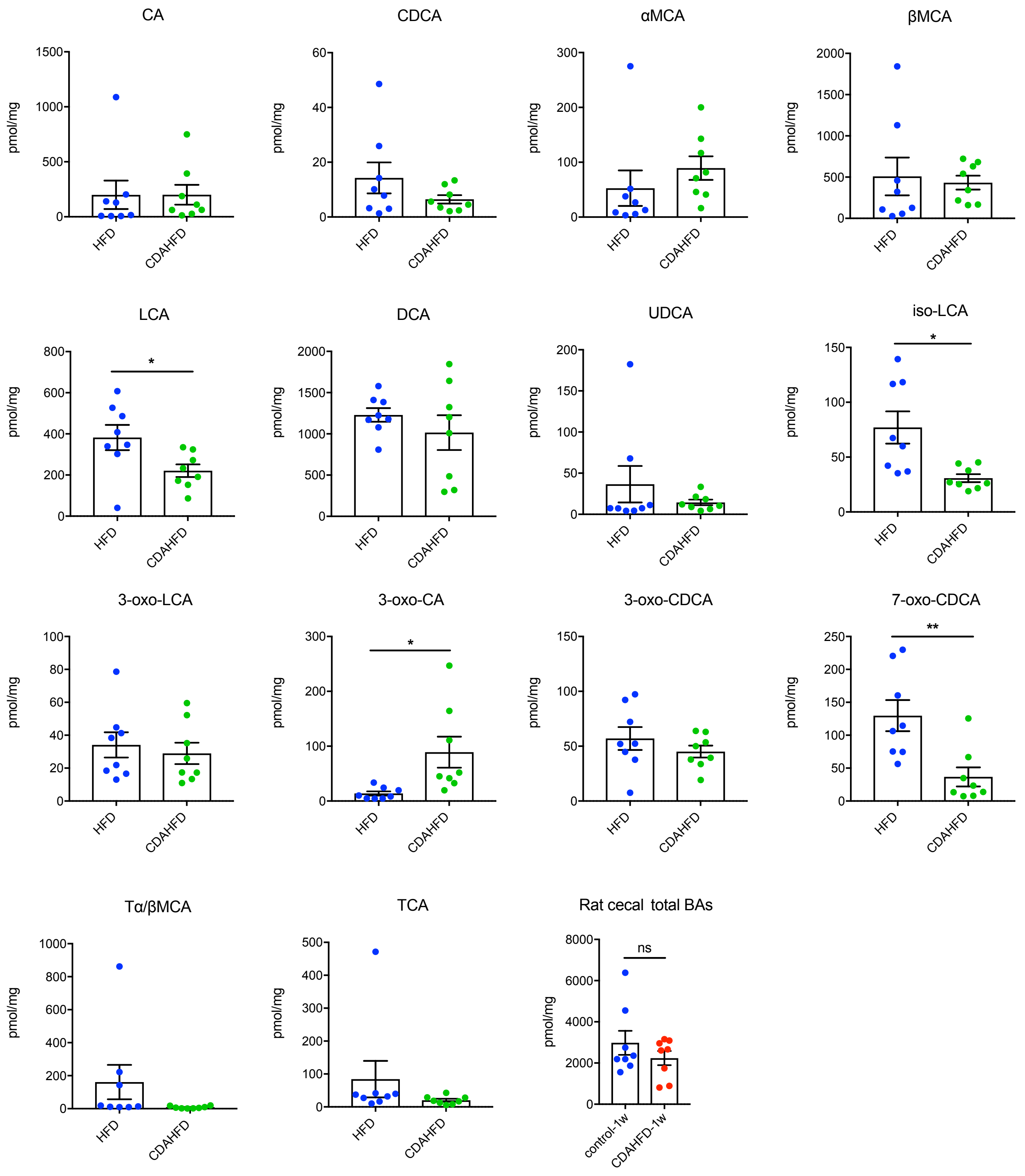

**Supplementary Figure 3**. **Bile acid concentrations in cecal contents of rats 1w post HFD control or CDAHFD diet intervention**. Bile acids were quantified using UPLC-MS. All bile acids with measurable concentrations above the limit of detection are shown (n=8 per group, two-tailed Welch's t test, data not marked with asterisk(s) were not significant). ns = not significant, *$p<0.05$, **$p<0.005$. Bars represent mean ± SEM.

### Figure S4

#### Rat portal BAs, 48h

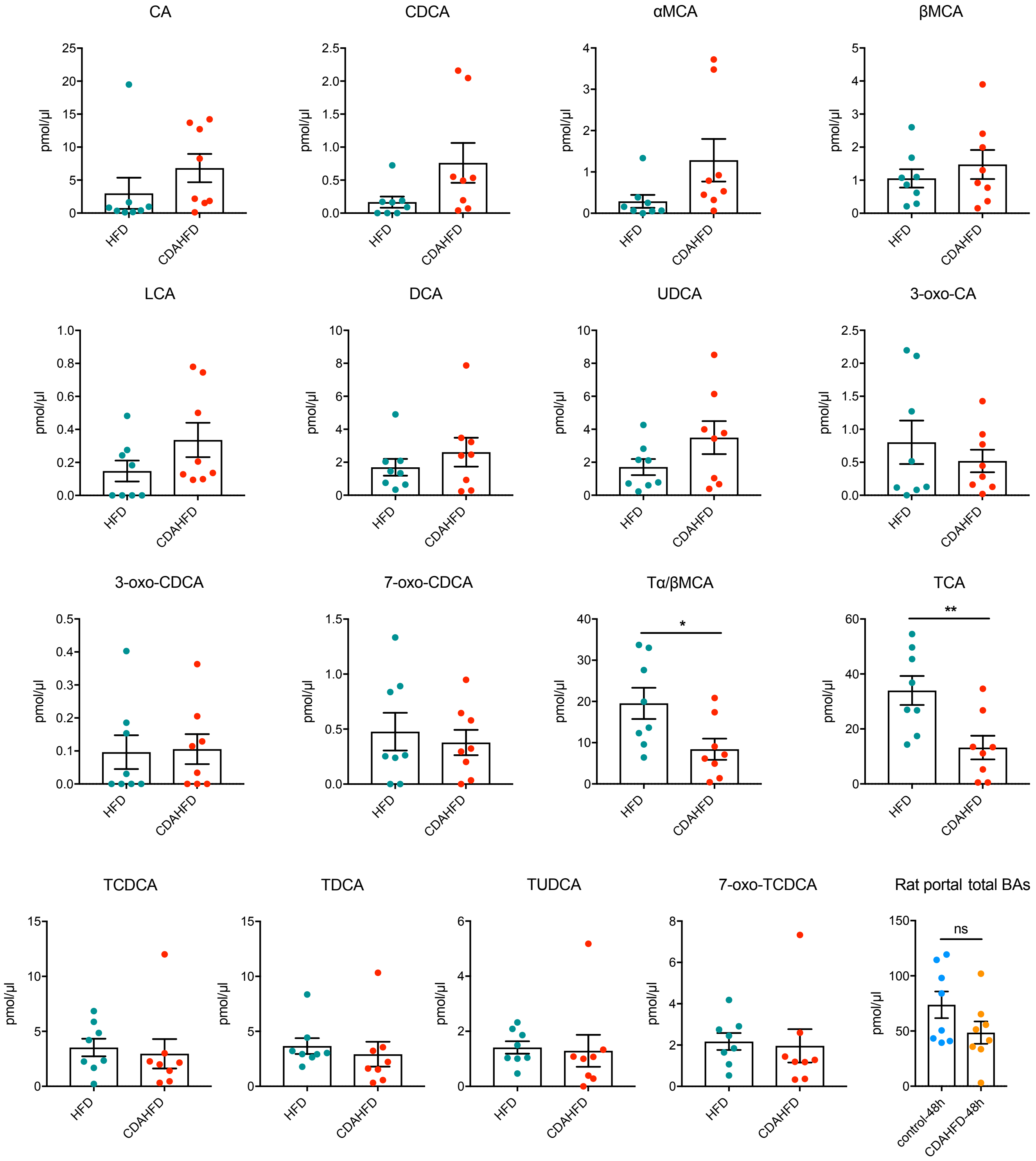

**Supplementary Figure 4**. **Bile acid concentrations in portal veins of rats 48h post HFD control or CDAHFD diet intervention**. Bile acids were quantified using UPLC-MS. All bile acids with measurable concentrations above the limit of detection are shown (n=8 per group, two-tailed Welch's t test, data not marked with asterisk(s) were not significant). ns = not significant, *$p$<0.05, **$p$<0.005. Bars represent mean ± SEM.

Figure S5

Rat portal BAs, 1w

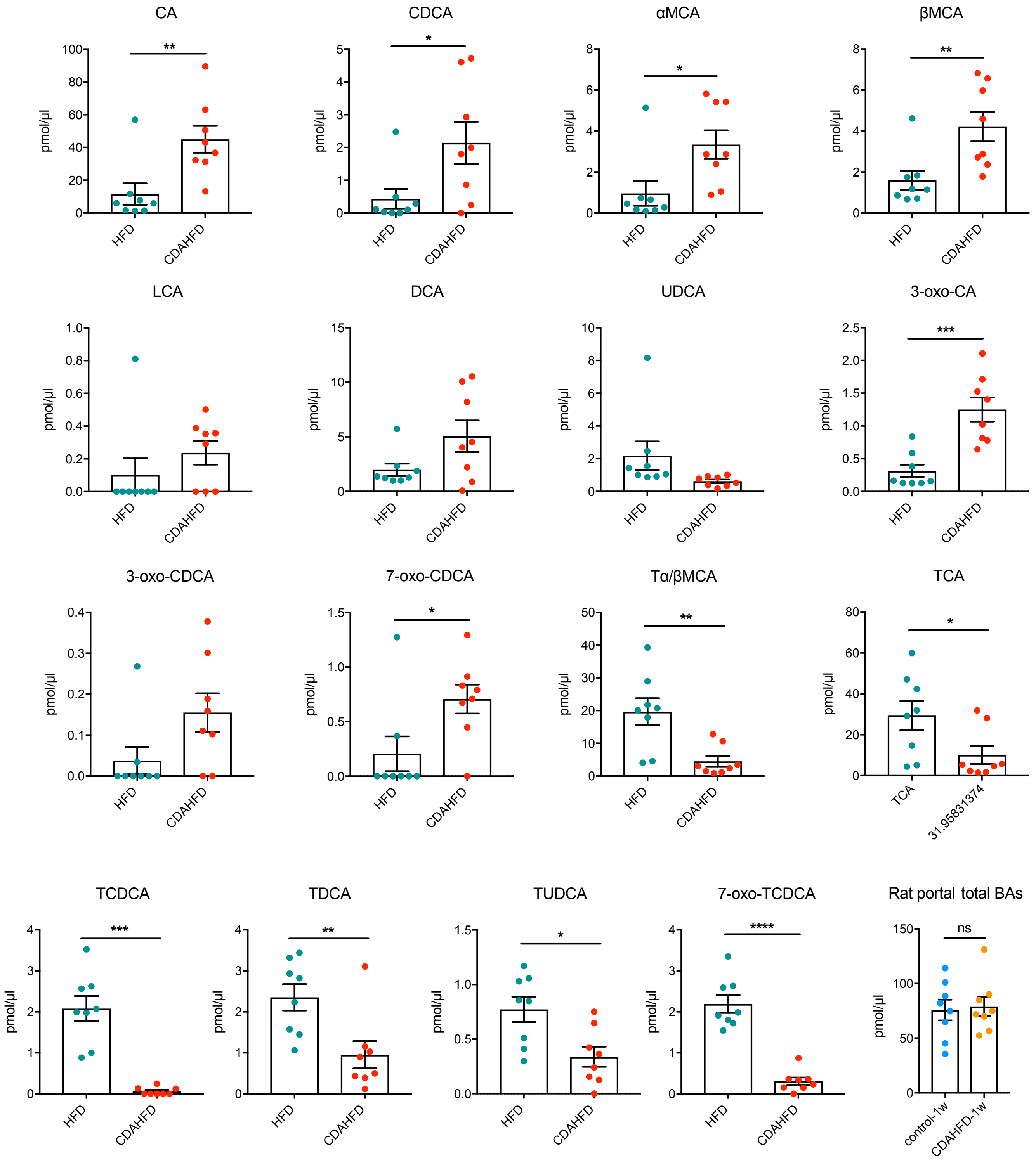

**Supplementary Figure 5**. **Bile acid concentrations in portal veins of rats 1w post HFD control or CDAHFD diet intervention**. Bile acids were quantified using UPLC-MS. All bile acids with measurable concentrations above the limit of detection are shown (n=8 per group, two-tailed Welch's t test, data not marked with asterisk(s) were not significant).

ns = not significant, *$p$<0.05, **$p$<0.005, ***$p$<0.001, ****$p$<0.0001. Bars represent mean ± SEM.

### Figure S6

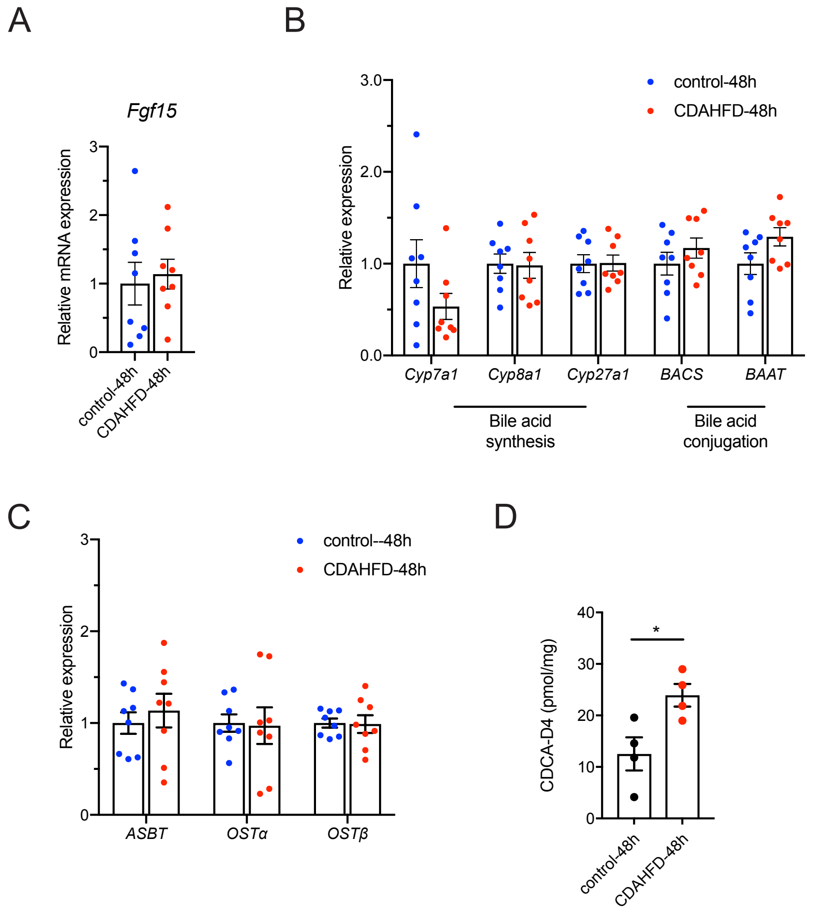

A
Fgf15
Relative mRNA expression
0
1
2
3
control-48h
CDAHFD-48h
B
Relative expression
0.0
1.0
2.0
3.0
control-48h
CDAHFD-48h
Cyp7a1
Cyp8a1
Cyp27a1
BACS
BAAT
Bile acid synthesis
Bile acid conjugation
C
Relative expression
0
1
2
3
control--48h
CDAHFD-48h
ASBT
OSTα
OSTβ
D
CDCA-D4 (pmol/mg)
0
10
20
30
40
*
control-48h
CDAHFD-48h

**Supplementary Figure 6**. **CDAHFD-fed rats display increased microbial BA deconjugation with no difference in FXR signaling, BA synthesis, conjugation, or transport.** (A) *Fgf15* expression as measured by RT-qPCR was unchanged 48 hours after dietary intervention (n=8 per group, two-tailed Welch's t-test)**.** (B, C) Liver mRNA expression of bile acid synthesis and conjugation genes (B) and intestinal BA transporters (C) measured by RT-qPCR was similar between control and CDAHFD-fed rats after 48h post-diet intervention (n=8 per group, two-tailed Welch's t test). (D) Cecal BSH activity was increased in CDAHFD-fed rats compared to control rats at 48h of diet as measured by conversion of deuterated glyco-CDCA (GCDCA-D4) to deuterated CDCA (CDCA-D4) (n=4 per group, two-tailed Welch's t test, data not marked with asterisk(s) were not significant). *$p<0.05$. Bars represent mean ± SEM.

Figure S7

Rat cecal BAs, 6w

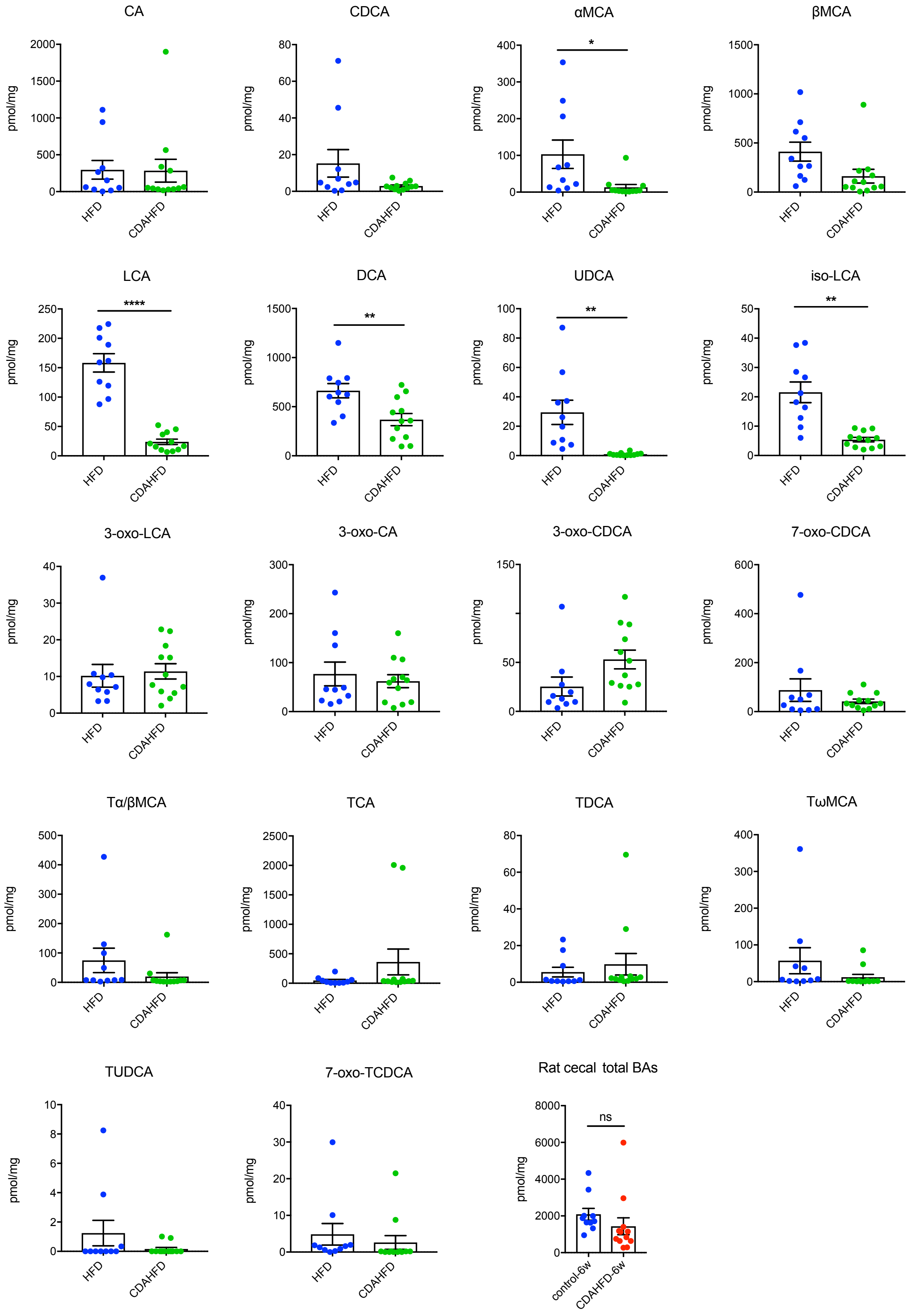

**Supplementary Figure 7**. **Bile acid concentrations in cecal contents of rats 6w post HFD control or CDAHFD diet intervention**. Bile acids were quantified using UPLC-MS. All bile acids with measurable concentrations above the limit of detection are shown (HFD n=10, CDAFD n=12, two-tailed Welch's t test, data not marked with asterisk(s) were not significant). ns = not significant, *$p<0.05$, **$p<0.005$, ****$p<0.0001$. Bars represent mean ± SEM.

Figure S8

#### Rat cecal BAs, 12w

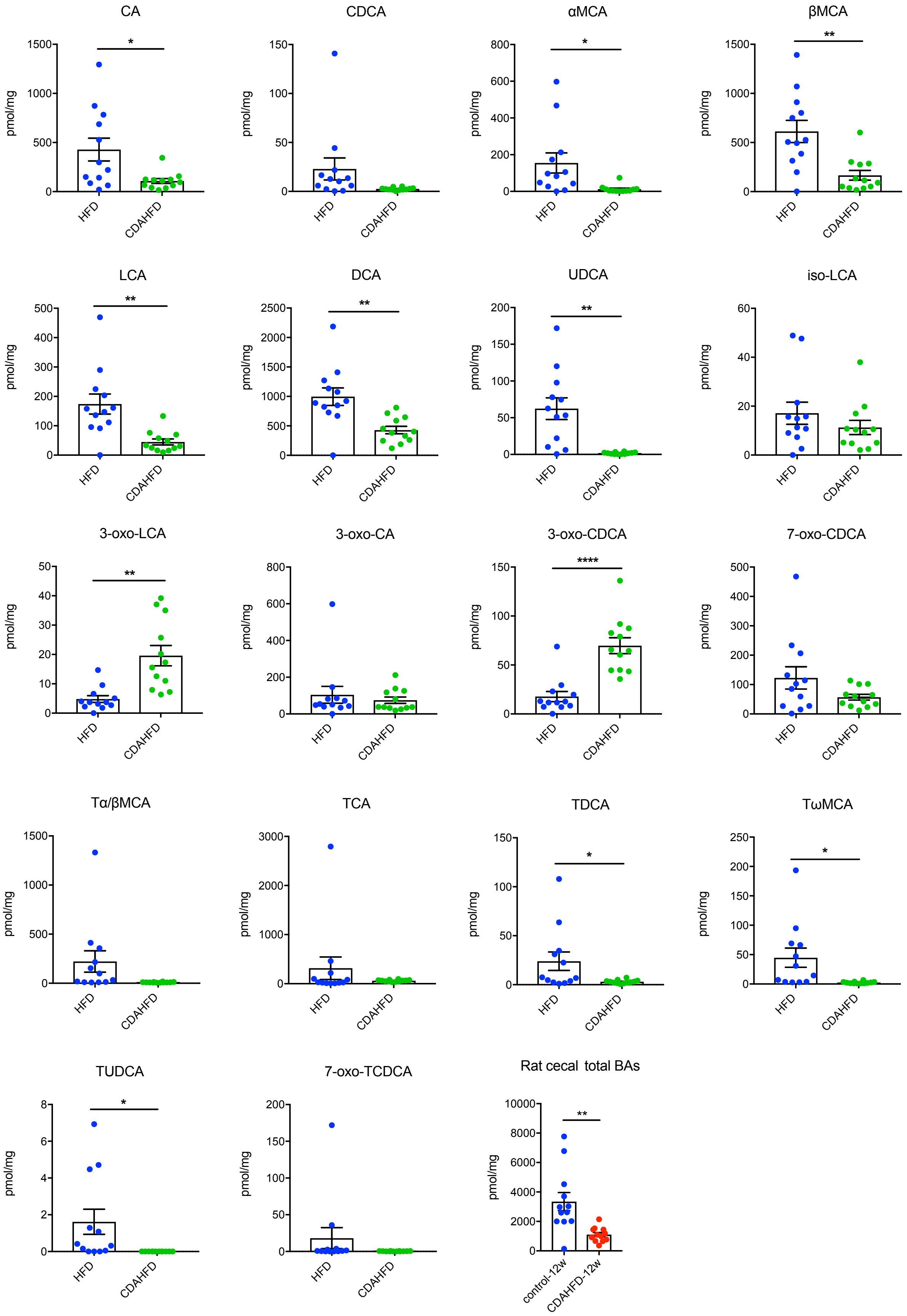

**Supplementary Figure 8**. **Bile acid concentrations in cecal contents of rats 12w post HFD control or CDAHFD diet intervention**. Bile acids were quantified using UPLC-MS. All bile acids with measurable concentrations above the limit of detection are shown (n=12 per group, two-tailed Welch's t test, data not marked with asterisk(s) were not significant). *$p<0.05$, **$p<0.005$, ****$p<0.0001$. Bars represent mean ± SEM.

### Figure S9

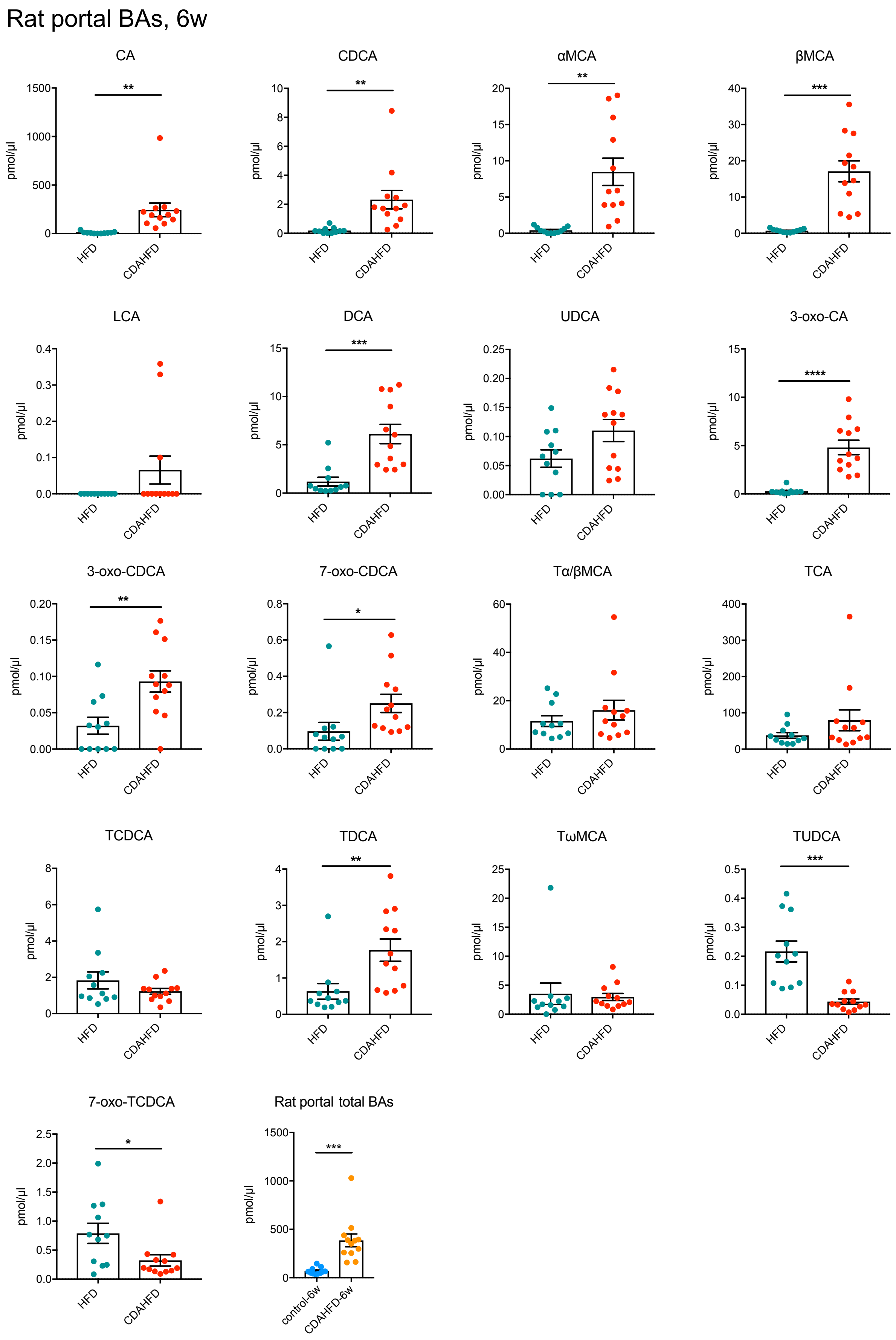

Rat portal BAs, 6w
CA
CDCA
αMCA
βMCA
LCA
DCA
UDCA
3-oxo-CA
3-oxo-CDCA
7-oxo-CDCA
Tα/βMCA
TCA
TCDCA
TDCA
TωMCA
TUDCA
7-oxo-TCDCA
Rat portal total BAs
pmol/μl
HFD
CDAHFD
control-6w
CDAHFD-6w

**Supplementary Figure 9**. **Bile acid concentrations in portal veins of rats 6w post HFD control or CDAHFD diet intervention**. Bile acids were quantified using UPLC-MS. All bile acids with measurable concentrations above the limit of detection are shown (n=12 per group, two-tailed Welch's t test, data not marked with asterisk(s) were not significant). *$p<0.05$, **$p<0.005$, ***$p<0.001$, ****$p<0.0001$. Bars represent mean ± SEM.

### Figure S10

#### Rat portal BAs, 12w

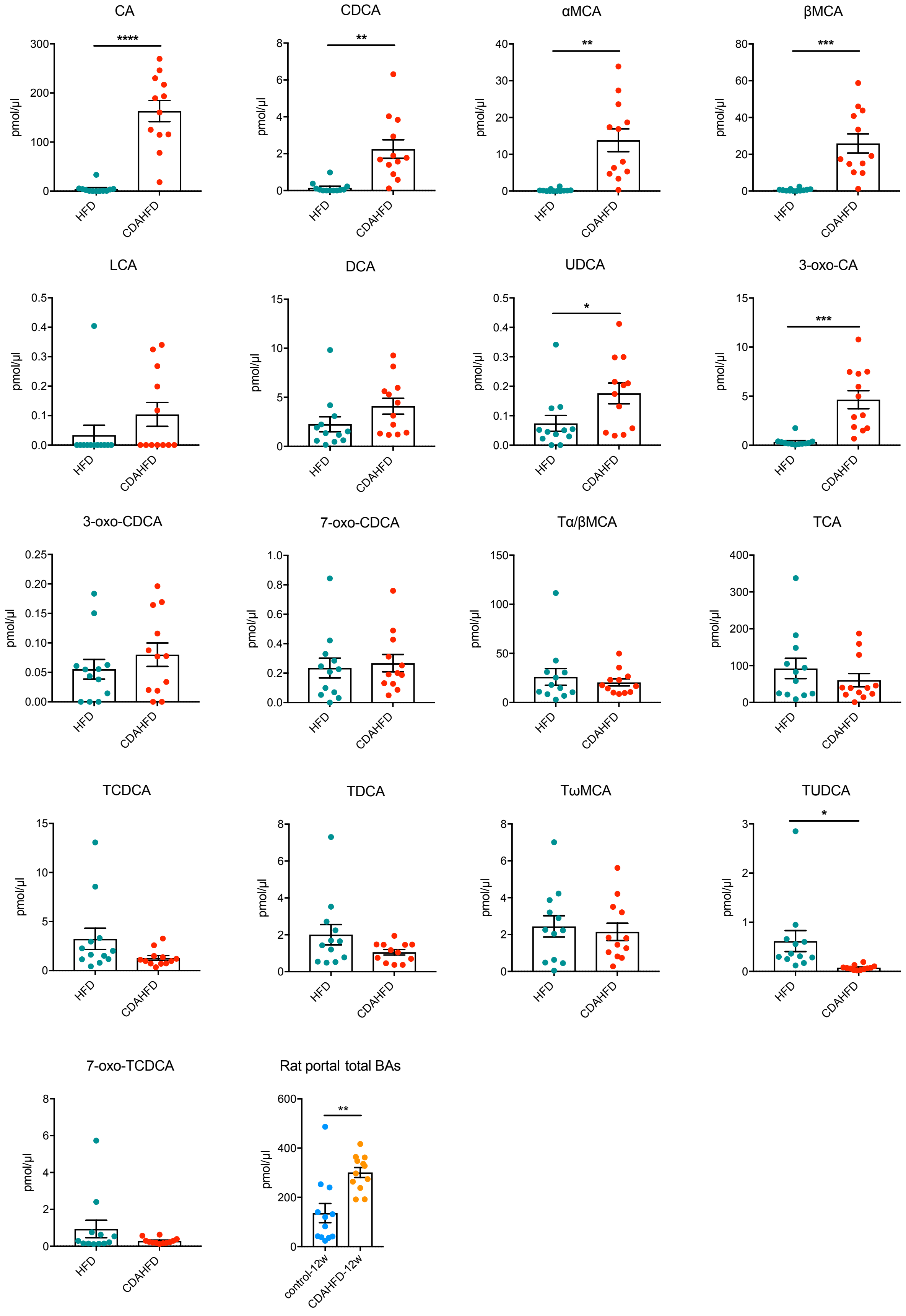

**Supplementary Figure 10**. **Bile acid concentrations in portal veins of rats 12w post HFD control or CDAHFD diet intervention**. Bile acids were quantified using UPLC-MS. All bile acids with measurable concentrations above the limit of detection are shown (n=12 per group, two-tailed Welch's t test, data not marked with asterisk(s) were not significant). **p*<0.05, ***p*<0.005, ****p*<0.001, *****p*<0.0001. Bars represent mean ± SEM.

Figure S11

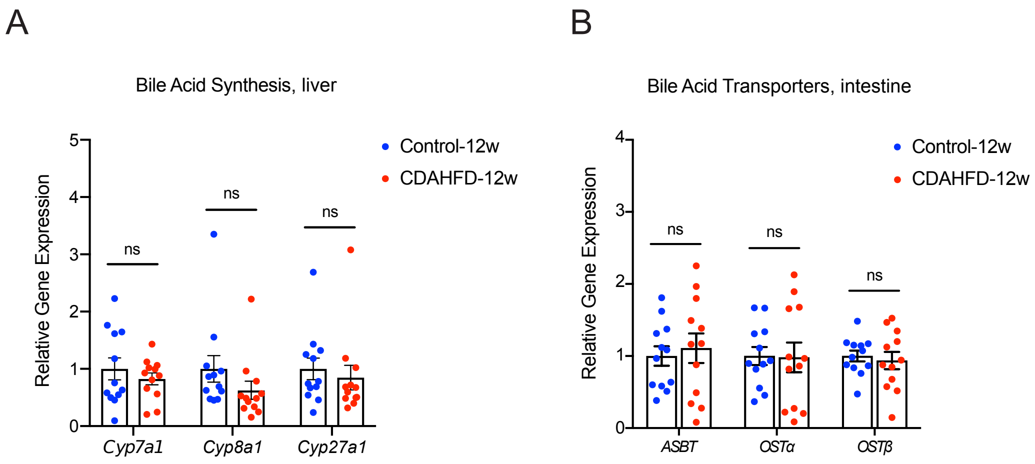
A
Bile Acid Synthesis, liver
Relative Gene Expression
0
1
2
3
4
5
ns
ns
ns
Cyp7a1
Cyp8a1
Cyp27a1
Control-12w
CDAHFD-12w
B
Bile Acid Transporters, intestine
Relative Gene Expression
0
1
2
3
4
ns
ns
ns
ASBT
OSTα
OSTβ
Control-12w
CDAHFD-12w

**Supplementary Figure 11**. **CDAHFD-fed rats at 12w show no difference in BA synthesis or transport.** (A, B) Liver mRNA expression of bile acid synthesis (A) and intestinal BA transporters (B) measured by RT-qPCR was similar between control and CDAHFD-fed rats after 12w post-diet intervention (n=12 per group, two-tailed Welch's t test). ns = not significant. Bars represent mean ± SEM.

Figure S12

Unconjugated BAs - 2mM

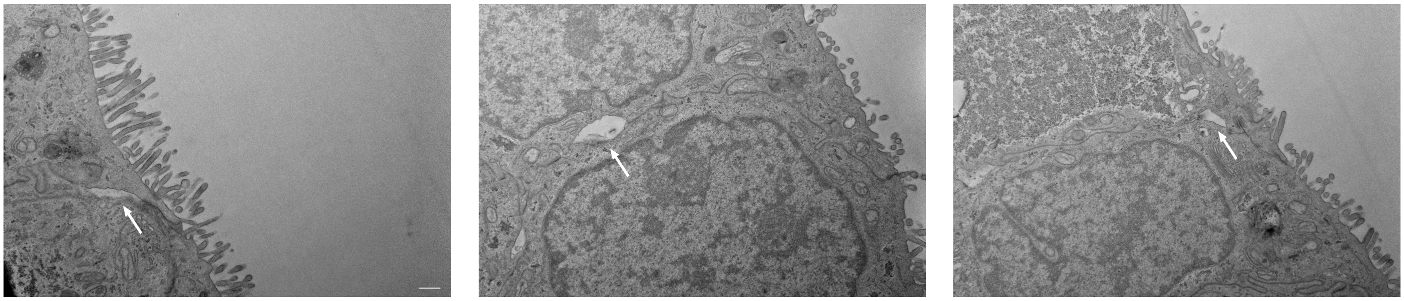

Conjugated BAs - 2mM

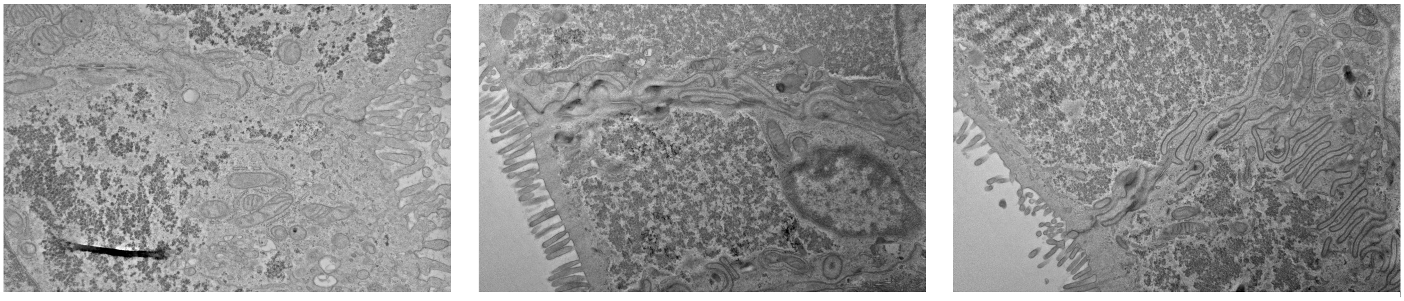

Unconjugated + conjugated - 2mM

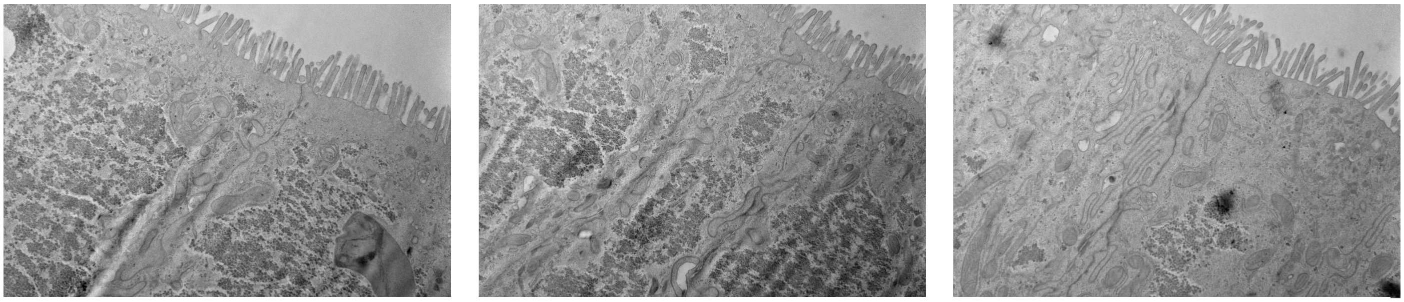

**Supplementary Figure 12. Conjugated BAs prevented the development of unconjugated BA-induced tight junction dilatation.** Additional representative EM images of Caco2 cells from transwells after exposure to conjugated, unconjugated, and combined BA pools at indicated concentrations. The white arrows point to examples of tight junction dilatation.

Figure S13

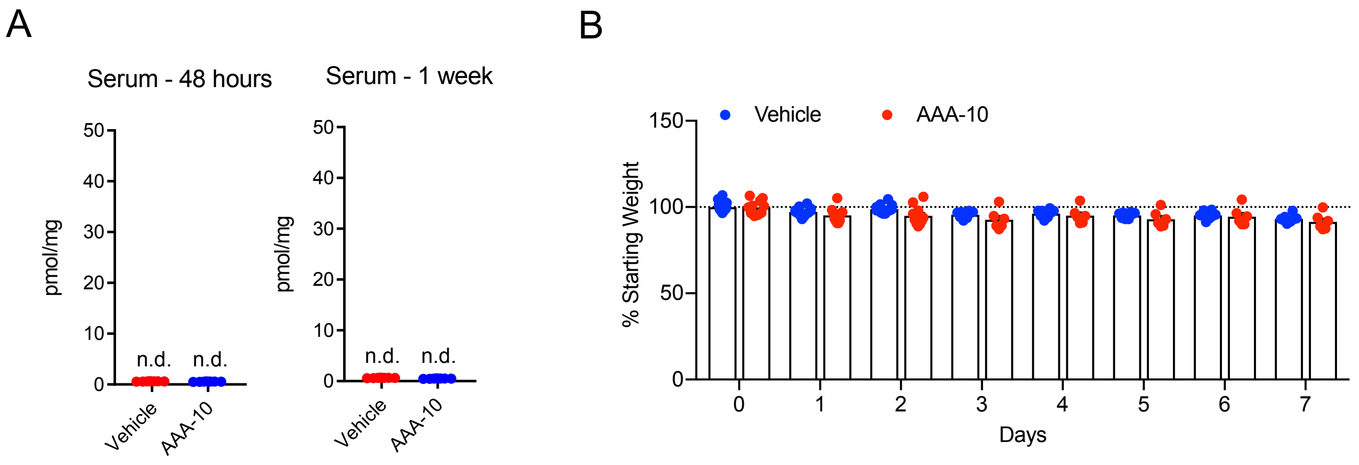
A
Serum - 48 hours
Serum - 1 week
pmol/mg
50
40
30
20
10
0
n.d.
n.d.
Vehicle
AAA-10
B
Vehicle
AAA-10
% Starting Weight
150
100
50
0
0
1
2
3
4
5
6
7
Days

**Supplemental Figure 13. Metrics for AAA-10 treatment of CDAHFD rats for 48h and 1w.** (A) UPLC-MS analysis of serum demonstrated no detection of AAA-10 in treated animals (n=8 per group, two-tailed Welch's t test). (B) Weight of vehicle and AAA-10 treated CDAHFD-fed rats over 7 days of treatment normalized to starting weight (n=8 per group, two-tailed Welch's t test, data not marked with asterisk(s) were not significant). ns = not significant. Bars represent mean ± SEM.

### 1w AAA-10 treatment, cecal BAs

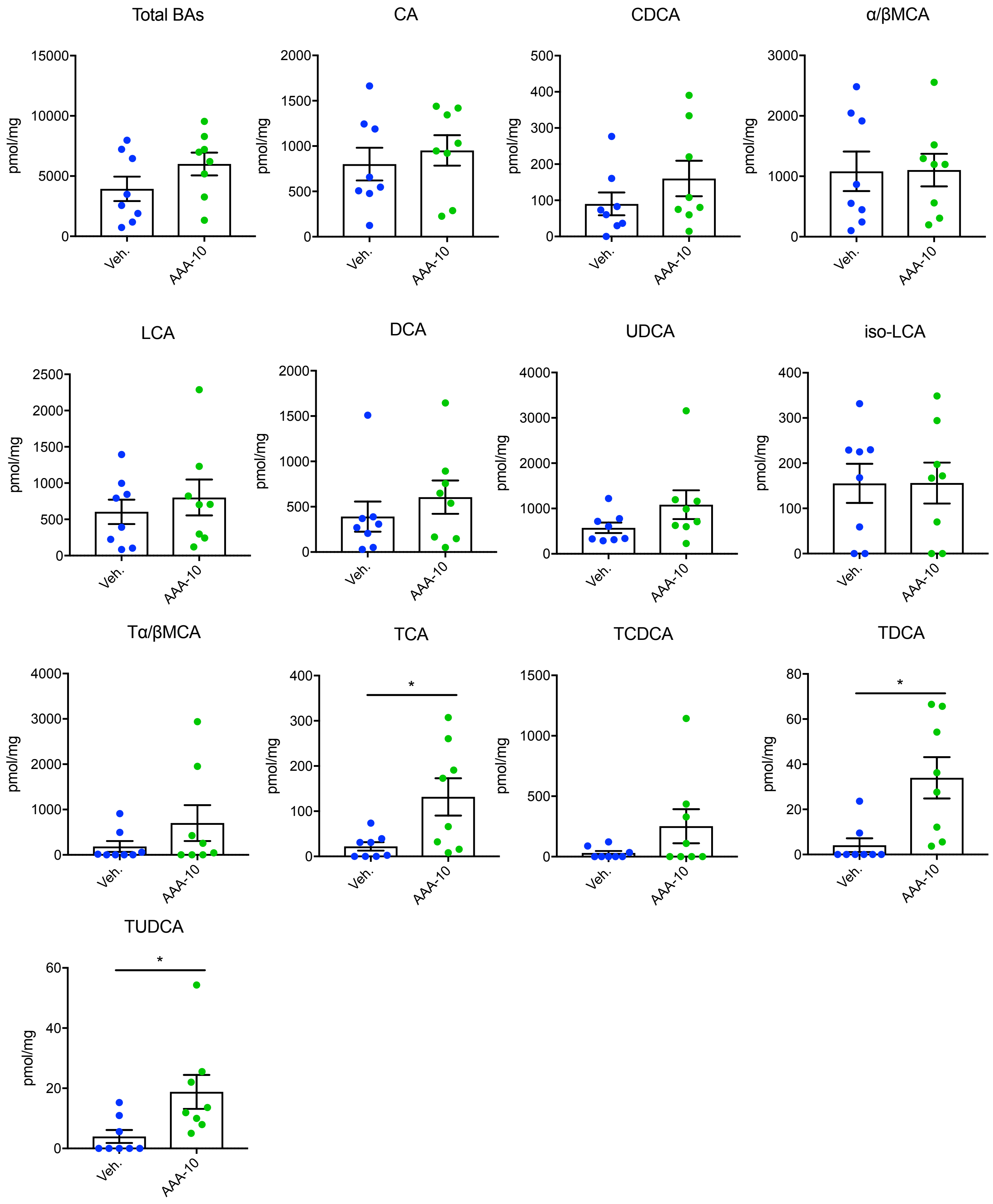

**Supplementary Figure 14**. **Bile acid concentrations in cecal contents of rats 1w post CDAHFD diet intervention and AAA-10 treatment**. Bile acids were quantified using UPLC-MS. All bile acids with measurable concentrations above the limit of detection are shown (n=8 per group, two-tailed Welch's t test, data not marked with asterisk(s) were not significant). *$p$<0.05. Bars represent mean ± SEM.

8 day AAA-10 treatment, cecal BAs

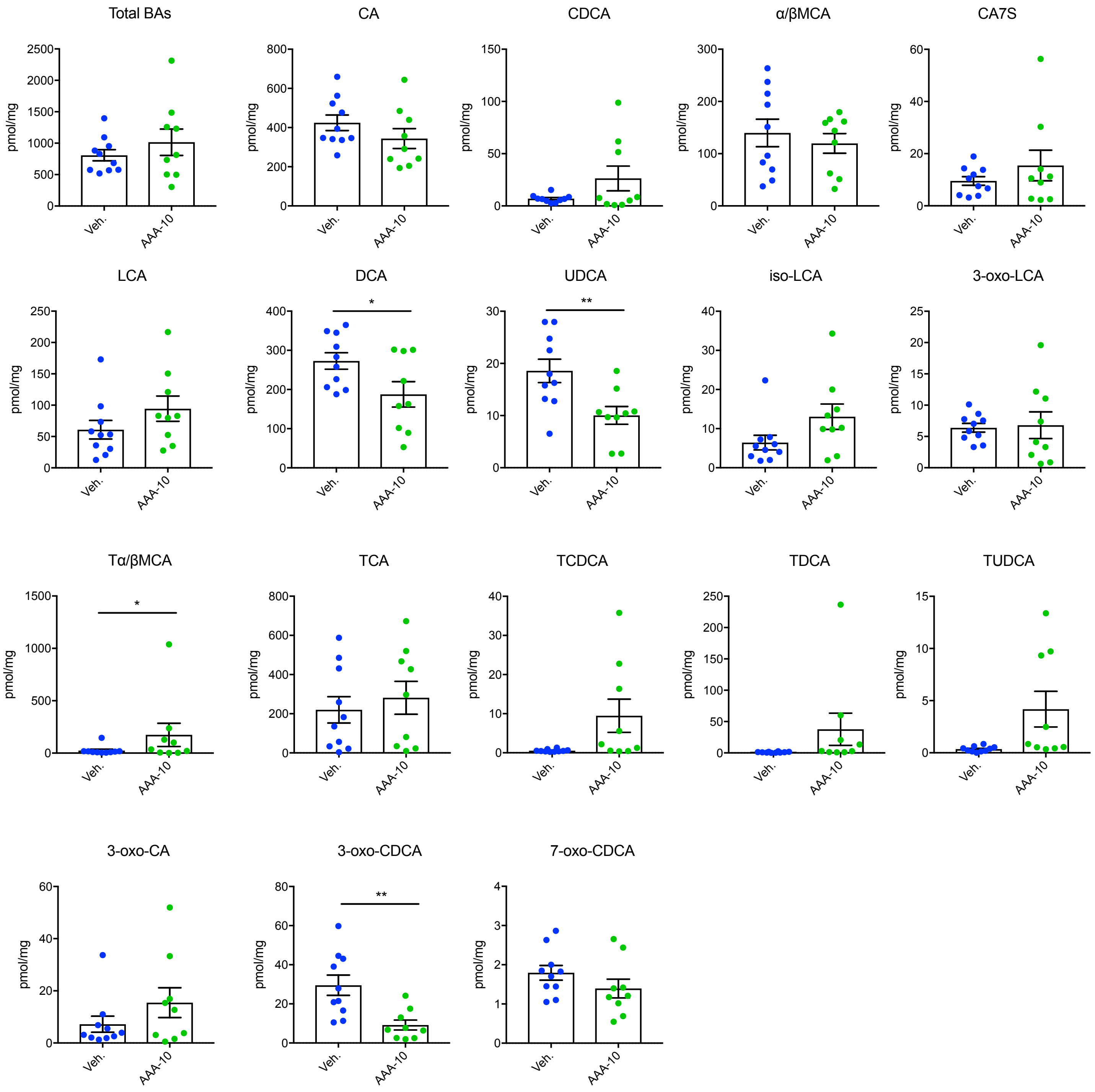

**Supplementary Figure 15**. **Bile acid concentrations in cecal contents of rats 8 days post CDAHFD diet intervention and AAA-10 treatment**. Bile acids were quantified using UPLC-MS. All bile acids with measurable concentrations above the limit of detection are shown (n=8 per group, two-tailed Welch's t test, data not marked with asterisk(s) were not significant). *$p<0.05$, **$p<0.005$. Bars represent mean ± SEM.

Figure S16

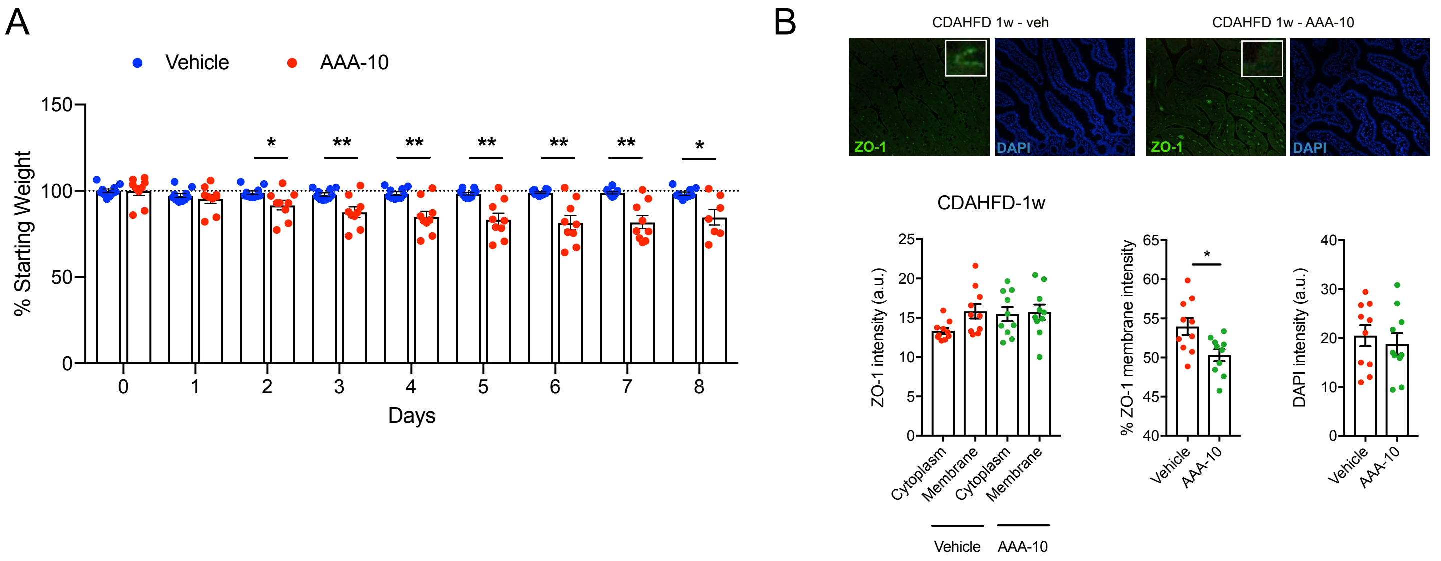

A
Vehicle
AAA-10
% Starting Weight
Days
B
CDAHFD 1w - veh
CDAHFD 1w - AAA-10
ZO-1
DAPI
CDAHFD-1w
ZO-1 intensity (a.u.)
Cytoplasm
Membrane
Vehicle
AAA-10
% ZO-1 membrane intensity
DAPI intensity (a.u.)

**Supplemental Figure 16. Metrics for AAA-10 treatment of CDAHFD rats for 8 days.** (A) Weight of vehicle and AAA-10 treated CDAHFD-fed rats over 8 days of treatment normalized to starting weight (n=10 in vehicle group, n=9 in AAA-10 group except day 8, where n=7 in AAA-10 group, two-tailed Welch's t test). (B) AAA-10 treatment prevented aberrant ZO-1 subcellular localization in absence of weight loss. ZO-1 immunofluorescence and DAPI counterstaining of rat ileum with quantification from vehicle and AAA-10 treated CDAHFD-fed rats at indicated timepoints. Intestines from 4 animals per group were stained (n=10 intestinal cell images per group were analyzed. For ZO-1 intensity, one-way ANOVA followed by Tukey's multiple comparison test, for %ZO-1 membrane intensity and DAPI intensity, two-tailed Welch's t test, data not marked with asterisk(s) were not significant). *$p<0.05$, **$p<0.005$. Bars represent mean ± SEM.

Figure S17

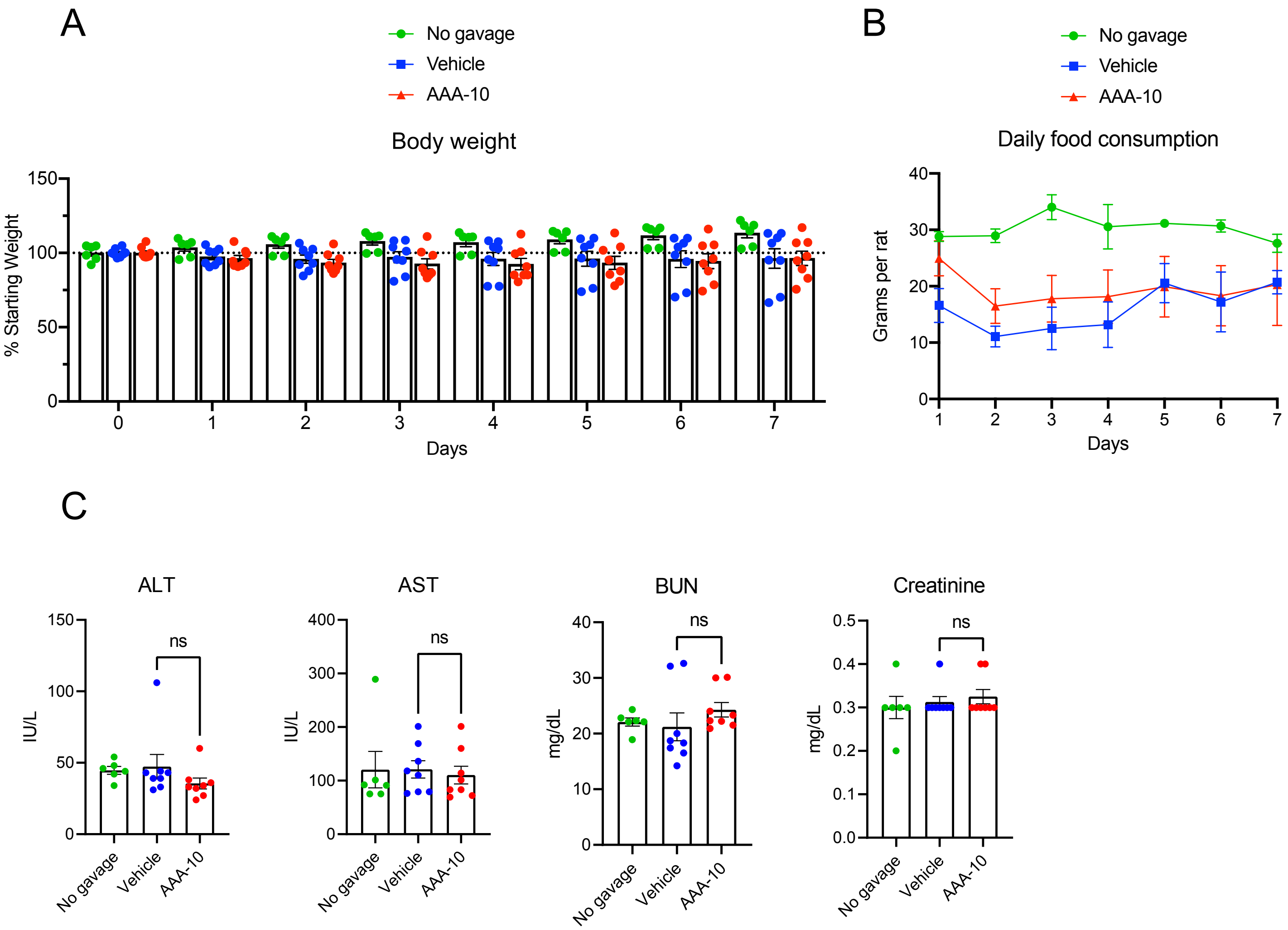
A
No gavage
Vehicle
AAA-10
Body weight
% Starting Weight
150
100
50
0
0
1
2
3
4
5
6
7
Days
B
No gavage
Vehicle
AAA-10
Daily food consumption
Grams per rat
40
30
20
10
0
1
2
3
4
5
6
7
Days
C
ALT
IU/L
150
100
50
0
ns
No gavage
Vehicle
AAA-10
AST
IU/L
400
300
200
100
0
ns
No gavage
Vehicle
AAA-10
BUN
mg/dL
40
30
20
10
0
ns
No gavage
Vehicle
AAA-10
Creatinine
mg/dL
0.5
0.4
0.3
0.2
0.1
0.0
ns
No gavage
Vehicle
AAA-10

**Supplemental Figure 17. Toxicity assessment of AAA-10 in chow-fed rats.** (A) No difference in body weight was observed between AAA-10-treated and vehicle-treated rats. Daily body weights measured and normalized to starting weight on day 0 (n=8 per group for both vehicle- and AAA-10 treated, n=6 in no gavage control, not significant, one-way ANOVA followed by Tukey's multiple comparison test). (B) AAA-10 did not induce differences in food consumption between vehicle and AAA-10 treated rats. Daily food consumption was monitored (n=8 per group for both vehicle- and AAA-10 treated, n=6 in no gavage control, not significant one-way ANOVA followed by Tukey's multiple comparison test). (C) Serum measurement of alanine aminotransferase (ALT), aspartate aminotransferase (AST), blood urea nitrogen (BUN), and creatinine (n=8 per group for both vehicle- and AAA-10 treated, n=6 in no gavage control, one-way ANOVA followed by Tukey's multiple comparison test, data not marked with asterisk(s) were not significant). ns = non-significant. Bars represent mean ± SEM.

**Supplementary Table 1.** Primer sequences used in this manuscript for quantitative PCR analysis.

| Gene | Forward Sequence (5' to 3') | Reverse Sequence (5' to 3') |
|---|---|---|
| *CYP7A1* | GGGCAGGCTTGGGAATTTTG | GGGCAGGCTTGGGAATTTTG |
| *CYP8B1* | CAGGTTGGAAGCCGAGACAT | CAGGTTGGAAGCCGAGACAT |
| *CYP27A1* | TGGACAACCACCTTTGGGAC | TGGACAACCACCTTTGGGAC |
| *BACS* | TTCAGGGACCACTGGACTTCCAAA | ACCACATCATCAGCTGTTCTCCCA |
| *BAAT* | GGTTGGCATCCTTTCTGTGTGCAT | ATTCTTCACTGCAGGGTGTAGGCT |
| *ASBT* | GGTTGCGCTTGTTATTCCTGT | GGTTCAATGATCCAGGCACTT |
| *OSTα* | GGCCCTTTCCAGTATGCCTT | CAGGTGCAACTTGGCTTGAC |
| *OSTβ* | GAAGCAGCCACAAGACAACG | TCTCTTAGGATGCCCAGGCT |
| *Col1a1* | TCTGACTGGAAGAGCGGAGA | GGGTTTGGGCTGATGTACCA |
| *Acta2* | GGAGATGGCGTGACTCACAA | CGCTCAGCAGTAGTCACGAA |
| *IL-1β* | CACCTCTCAAGCAGAGCACA | ACGGGTTCCATGGTGAAGTC |
| *Tnfα* | ATGGGCTCCCTCTCATCAGT | TTTGCTACGACGTGGGCTAC |
| *IFNγ* | CGAGGTGAACAACCCACAGA | TTTGCTACGACGTGGGCTAC |
| *Cxcl2* | ACCATCAGGGTACAGGGGTT | CAACCCTTGGTAGGGTCGTC |
| *Cxcl9* | GTTTGCCCCAAGCCCTAACT | GCTGAATCTGGGTCTAGGCA |
| *Cxcl16* | TTTGGACCCTTGGCCCTTAC | AGTAGCAACTTCCAGCGACA |
| *Ccl2* | AGTTAATGCCCCACTCACCTG | GTAGTTCTCCAGCCGACTCA |
| *GAPDH* | ATGACTCTACCCACGGCAAG | CTGGAAGATGGTGATGGGTT |
